# Single-Cell Classification Using Graph Convolutional Networks

**DOI:** 10.1101/2021.06.13.448259

**Authors:** Tianyu Wang, Jun Bai, Sheida Nabavi

**Author notes:** **Corresponding author:** Sheida Nabavi.

## Abstract

**Background:** Analyzing single-cell RNA sequencing (scRNAseq) data plays an important role in understanding the intrinsic and extrinsic cellular processes in biological and biomedical research. One significant effort in this area is the identification of cell types. With the availability of a huge amount of single cell sequencing data and discovering more and more cell types, classifying cells into known cell types has become a priority nowadays. Several methods have been introduced to classify cells utilizing gene expression data. However, incorporating biological gene interaction networks has been proved valuable in cell classification procedures.

**Results:** In this study, we propose a multimodal end-to-end deep learning model, named sigGCN, for cell classification that combines a graph convolutional network (GCN) and a neural network to exploit gene interaction networks. We used standard classification metrics to evaluate the performance of the proposed method on the within-dataset classification and the cross-dataset classification. We compared the performance of the proposed method with those of the existing cell classification tools and traditional machine learning classification methods.

**Conclusions:** Results indicate that the proposed method outperforms other commonly used methods in terms of classification accuracy and F1 scores. This study shows that the integration of prior knowledge about gene interactions with gene expressions using GCN methodologies can extract effective features improving the performance of cell classification.

## Background

With the explosion of single cell RNA sequencing (scRNAseq) technology in recent years, the unprecedented opportunity for single cell transcription analysis has emerged. The traditional bulk RNA sequencing methods sequence a mix of millions of cells. This results in gene expression of a gene that reflects an average value of the gene expression across all the cells, overlooking the heterogeneity between cells. Different from bulk RNAseq, scRNAseq isolates cells in the first step and performs sequencing on thousands of genes for each cell in the second step. According to different sequencing protocols, hundreds to millions of expression values are gathered for each gene, which allows to identify new cell types [1–3], identify gene regulatory mechanisms, and solve the cellular dynamics of developmental processes.

Data analysis for scRNAseq data has attracted much research efforts, such as differential expression analysis, cell clustering, and missing value imputation. In early studies, most researchers have exploited the clustering analysis to categorize cell types [4, 5]. Since most cell sub-types have been identified and features of transcriptomics have become available, the focus on identifying cell subtypes has shifted from detecting new cell types based on clustering-based methods to discovering cell-specific expression signatures based on classification-based methods. Classification is a supervised learning method that requires plenty of data to perform the training process. Compared to clustering, a well-trained classifier based on public annotated datasets can efficiently and accurately identify the unlabeled cells, even the unlabeled data are from different platforms and samples [6].

Few cell classification methods have been proposed for scRNAseq data that can be grouped into two categories. One group of cell classification methods, including scPred [7], scID [8] and CasTLe [9] are based on traditional supervised machine learning methods, such as random forest (RF), support vector machine (SVM) and k nearest neighbors (KNN) [7, 10]. scPred [7] combines a feature selection and SVM with radial kernel to classify cell types. scID [8] applies the fisher linear discriminant analysis for cell classification. CasTLe [9] combines the data preprocessing and classifies cells based on the XGBoost [11] classifier. The other group of methods are based on cluster level similarity measurements, for example SingleR [12] and scmap [13]. SingleR [12] employs Spearman correlation to calculate the similarity between cells. There are two methods in the scmap [13] package, scmapcluster and scmapcell. scmapcluster, first, constructs a virtual cell for each cell type based on the median expression values of the genes across that specific cell type. Then, it computes the cell similarity between the cells that need to be labeled and the virtual cells. The labels are assigned by the highest similarity. Scmapcell computes the cell similarity directly and employs the KNN method to classify the cells. Also, some studies have used marker genes as prior knowledge for cell classification [14, 15]. However, the marker genes are not always available. These classification methods are not scalable and they work well only when dealing with small datasets with the prerequisite of reasonable feature selection.

With the huge progress of deep learning in computer vision and image classification, a variety of deep learning models have been explored and designed. A deep learning network learns high-level features from data and thus does not need the domain knowledge to select the features, which is beneficial for the classification of a huge number of samples. A recent cell classification method, ACTINN [16], employs a fully connected neural network for cell type classification.

Even though gene expression values are regulated by gene networks, classification based on gene expression values ignores this prior gene interaction knowledge. To combine the prior knowledge of biological networks and gene expression, we introduce the use of graph convolutional network (GCN) [17, 18]. The gene network and gene expression can be considered as different views of the data. To learn useful features to represent complex data, multiple views of data need to be considered. This has motivated us to develop a multimodal deep learning model based on GCN and neural network (NN) methodologies. A GCN is an extension of convolution on the graph domain. The GCN approaches have been applied to address biological problems such as predicting protein functions [19, 20].

For single cell classification, this paper proposes a new GCN-based end-to-end multimodal deep learning model. The proposed model learns the feature map based on both the gene expression values and the gene-gene interaction structure. The proposed multimodal GCN model employs a GCN paralleled with an NN model. Due to the localization property of learning filters in the GCN, it can extract local features based on the prior knowledge – the genomics interaction network. Hence, employing only a GCN model fails to capture global features. The quality of the extracted features depends on not only the GCN filters but also the completeness of the underlying genomics interaction network. The features extracted by a fully connected NN can represent the global connections but neglect the inner interactions. As a result, to have a more precise feature embedding, we utilize a parallel structure that consists of a GCN and an NN. In other words, combining localized features extracted from a GCN and global features extracted from an NN for classification conquers the limitations of GCN and NN models. We evaluate the performance of the proposed method using seven single cell datasets. We compare the performance of the proposed method with those of seven scRNAseq data classification tools and four conventional classification methods.

## Methods

### Network structure

The overall structure of the proposed model is shown in Figure 1. The proposed classification model consists of two parallel networks: a GCN and an NN. Gene expression values and the gene adjacency network are the inputs of the GCN; while the gene expression values are the inputs of the NN. The features learned by the GCN and the NN are concatenated and then connected with a fully connected layer. The output layer outputs the predicted cell type of an input cell.

**Figure 1:**
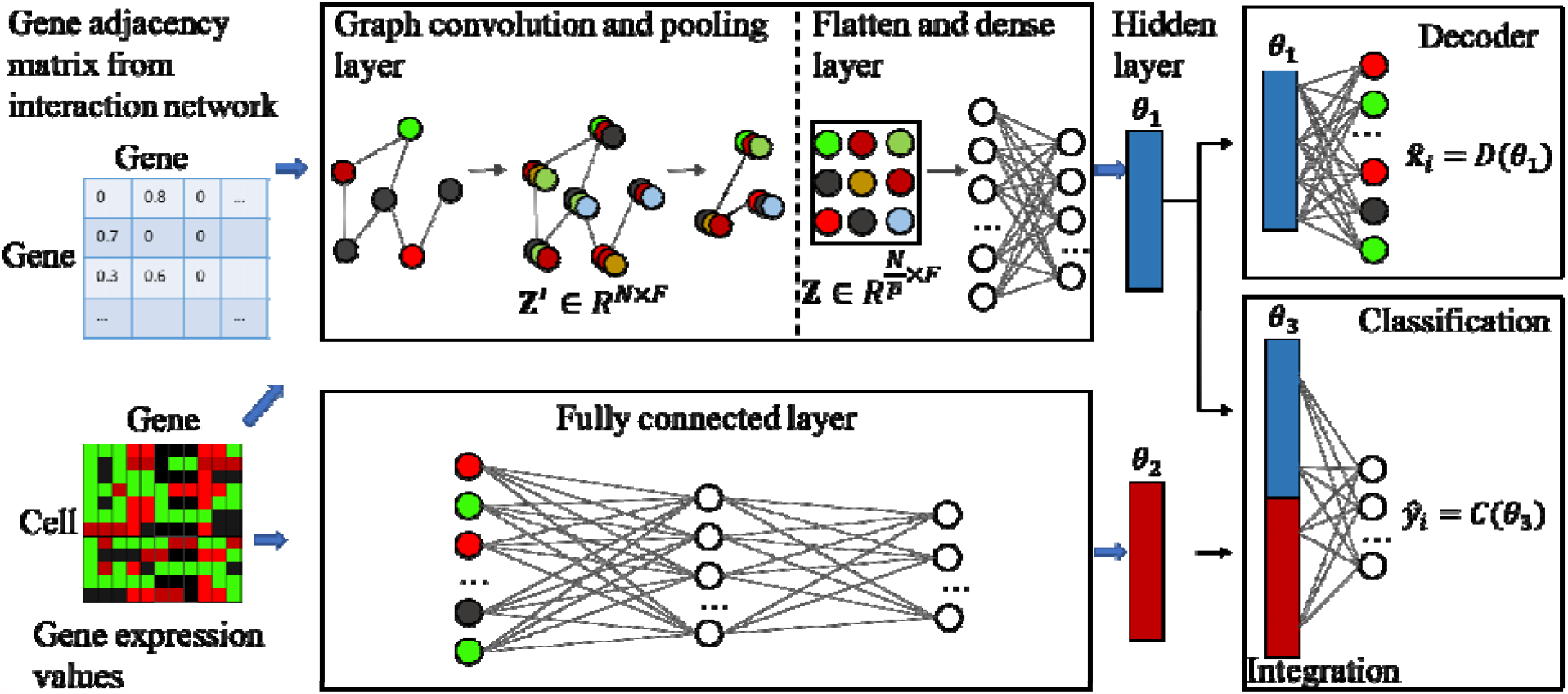
Structure of the proposed deep learning network (sigGCN) for single cell classification.

### Gene adjacency matrix

We utilized the STRING database [21] to build the weighted gene adjacency matrix. The size of the adjacency matrix is *N*× *N*, where *N* is the number of the genes. The elements in the matrix represent the confident score between pairs of genes extracted from the gene-gene interaction database. Then, we normalized the weights in the adjacency matrix by row sums. Using the normalized adjacency matrix, we built a weighted graph that the nodes are genes (proteins), the edges represent the connection between genes and the normalized confidence scores are weights of edges. We regarded the genes that have no neighbors in the matrix as the singletons in the network. We need to mention that since we chose top *N* genes with highest variances in expression values for training (explained in the Data preprocessing section), there are singletons in the graph. We discussed the model performance including the singletons and excluding the singletons in the results section.

To explore the benefit of using the gene network, we also constructed the gene adjacency matrix using the gene co-expression similarity, where the elements in the gene adjacency matrix are the gene cosine similarities. To compute the cosine similarity between two genes, we considered the gene expressions across all the cells as vectors. Then the cosine similarity of a pair of genes were computed by (**x**·**y**)/(∥**x** ∥∥**y**∥), where **x** and **y** are the expression vectors of the two genes. We filtered out edges for pairs of genes whose cosine similarities are less than 0.6. We compared the model performance when using the gene interaction network with that of when using the gene co-expressions to construct the gene graph.

### Graph convolutional network

In this work, to extract the features of expression values incorporating the gene interaction network, we developed a GCN-based autoencoder model. For the GCN analysis, gene expression values are assigned as the features of the nodes. The encoder part of the GCN model consists of a GCN layer followed by a maxpooling layer, a flatten layer and a fully connected (FC) layer. While the decoder part consists of a FC layer to reconstruct the gene expression values.

The GCN layer consists of graph convolution and pooling operations. The gene expression matrix can be represented as **X** ∈ *R*^*N⨯M*^, where *N* is the number of genes and *M* is the number of cells. Considering graph *G* = (*V,E)*, where *V* represent the vertices (genes) and *E* represents the edges between the vertices, the gene expression values can be regarded as the vertex features. The adjacency matrix **A** ∈ *R*^*N⨯N*^ is used to represent the edges, i.e. the connections between the genes, constructed from the gene-gene interaction network. The Laplacian matrix is defined as **L** = **D** − **A**, where **D** ∈ *R*^*N⨯N*^ is a diagonal matrix. In the diagonal matrix D, the values on the diagonal represent the number of edges that connect to a vertex. The Laplacian matrix is further normalized to **L** = **I** + **D**^-1/2^**AD**^1/2^, where **I** ∈ *R*^*N⨯N*^ is the identity matrix. Because the normalized Laplacian matrix is a real symmetric positive semidefinite matrix, it can be factorized to:

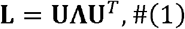

where **U** =(**u**_1_,…**u**_*l*_, …**u**_*n*_) contains a set of orthonormal eigenvectors, **UU**^T^ = **I**, and **Λ** is the eigenvalue matrix **Λ** = *diag*(*λ*_l_,… *λ*_*l*_,… *λ*_*n*_).

The graph convolutional theory comes from the spectral convolution theory. Like the convolution of two one-dimensional signals, given the Fourier transformation of a graph as 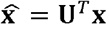 [22] and the inverse Fourier transformation 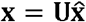, the convolution on the graph is defined as [18]:

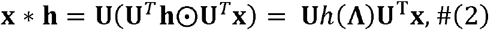

where ⊙ is the Hadamard (element-wise) product, and **x** represents the *N*−dimensional vector on the graph, i.e. the gene expression values of a cell in this work. *H*(**Λ**) is a diagonal matrix that is denoted as the convolution kernel of the transformation. In general, the designation of the convolution kernel decides the computation cost of the graph convolution. The convolution kernel can be designed as 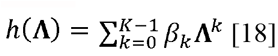, where *K* is the receptive field of the convolution kernel (i.e. the order of the neighbors that are computed) and *β*_*k*_s are the polynomial coefficients. The kernel is approximated by the Chebyshev polynomial as following to further decrease the computation cost:

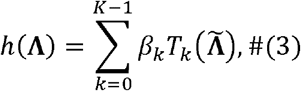

where 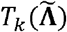 is the *k*^*th*^ order of the Chebyshev polynomial which is recursively defined as 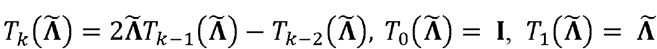 [23]. The 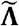 is the rescaled diagonal eigenvalue matrix of the Laplacian matrix, which is defined as 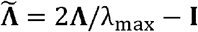. The parameters, *β*_*k*_s, are learned during the training process. Therefore, when substituting Equation (3) into Equation (2), the convolution on the graph is computed by:

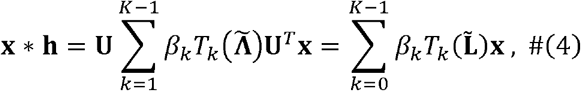

where 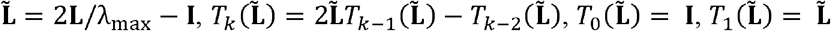. With this substitution, the multiplication of the matrices is no longer needed, and the convolution computation depends on the *K*^th^ order neighbors of the vertices. In this way, the convolution is transformed to the weighted summation of the *K*-hop neighbors which can keep the spatial localization. Then, the output of the graph convolution layer can be written as:

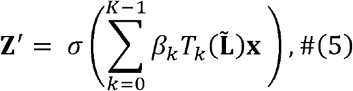

where *σ*(·) is the activation function, and **x** is the input of the graph, i.e. the gene expression values of a cell. The receptive field of the kernel, *K*, can be considered as a hyper-parameter which is chosen as 5 in this study. In this work, we employed multiple convolution kernels (*F* = 5), then the dimension of the graph convolutional layer output feature map is *N* × *F*. We also used a maxpooling layer with size *p* = 8 after the graph convolutional layer, which means we clustered *p* nodes (genes) in the graph into one big node. Therefore, the feature map (**Z**) generated from the proposed GCN has the dimension of the number of nodes after pooling (*N*/*p*) by the number of features (F = 5), Z ∈ *R*^*N/PXF*^.

Since the main goal is classification on the graph level (cell level) but not on the node level (genes), this feature map is flattened and then connected with a dense layer with the size of 32 neurons to reduce the dimension. Therefore, the final feature, *θ*_1_(shown in Figure 1), outputted by the GCN is a vector with the size of 32.

### Reconstruction of gene expression

In the GCN-based autoencoder, the decoder part is utilized to reconstruct the node attributes (gene expressions). The features learned by the GCN (encoder part), i.e. the hidden layer shown in Figure 1, is fully connected to an output layer to build the decoder network. The output of the decoder has the same dimension as the input, which is the number of input genes, *N*. We utilized the mean squared error (MSE) as the loss function for this task:

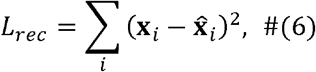

where **x**_*i*_ is the vector of gene expression values of cell *i* and 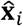 is the vector of reconstructed gene expression values.

### Neural network

The inputs of the NN (Figure 1) are the gene expression values of the top *N*(default is 1000) variant genes. Since the number of cells varies in different datasets and not all datasets have a large number of cells, we designed a shallow NN to avoid the overfitting problem. The network has two fully connected layers with the size of 256 and 32 neurons, respectively. The output of each layer is activated by the ReLU function. Therefore, the final feature from the NN, *θ*_2_(shown in Figure 1), is of size 32.

### Fully connected integrative layer for classification

The features learned by the NN are concatenated with the features learned by the GCN in the integration layer *θ*_3_(shown in Figure 1). The concatenated features are the input to the final output classification layer with the size of the number of the classes. Assume that there are *n* classes of cell types, the output layer is of size *n* and the corresponding probability array, **p** = [*p*_1_ … *p*_*n*_], is calculated over the *n* output neurons using the softmax activation function. The predicted label of cell *i* is the class that has the highest probability in array p, which is ŷ_*i*_ =argmax (**p**), shown in Figure 1. For its corresponding true class *y*_*i*_, the loss is computed by the negative log-likelihood function defined as:

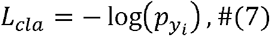

where 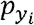 is the output probability of class (cell type) *y*_*i*_.

### Loss functions and model training

We also used the regularization loss *L*_*reg*_ to regularize the parameters and prevent overfitting. Assume W consists of all the parameters in the model — including the parameters in the GCN, the NN, the decoder, and the integration layer — and *w*_*j*_ represents each of the parameters in the model, the regularization loss is defined as:

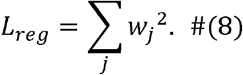

The total loss is a combination of the loss of classification, the loss of the reconstruction, and the loss of regularization:

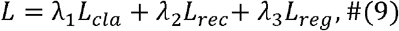

where *λ*_l_, *λ*_2_ and *λ*_3_ are the weights for each loss function. In this study, *λ*_l_ and *λ*_2_ are set to 1, and *λ*_3_ is set to 0.0001. The loss is computed and backpropagated to update the parameters. We used the SGD optimizer [24] to train the end-to-end model. The mini-batch size and epoch were chosen as 64 and 50, respectively. To train the model, we split the datasets into 80% as the training dataset, 10% as the validation dataset, and 10% as the testing dataset. We first used training dataset to train the model, and then tested the model on the validation dataset. We tuned the hyperparameters based on the performance of the model on the validation dataset. After tuning, we fixed the hyperparameters and evaluated the performance of the model on the testing dataset.

### Datasets

In this section, we describe the datasets and also the data preprocessing used in this work.

### Single cell datasets

To evaluate the performance of the proposed classification method, we used seven datasets which are from different sequencing protocols, across different species: human and mouse. All the datasets can be downloaded from https://doi.org/10.5281/zenodo.3357167 [25]. The datasets are across different cell populations and have a different number of cells and genes. Two of them are large scale datasets (Zheng68K and Zhengsorted) containing several ten-thousands of cells, and only one of the dataset (Zhengsorted) has validated ground truth while others use labels obtained by employing a clustering method on some marker genes.

#### Pancreatic datasets

We used five scRNAseq datasets from the human pancreas and mouse pancreas. The datasets are from different individual samples using different sequencing protocols. The BaronMouse [26] and the BaronHuman [26] datasets are from the mouse and human pancreas, respectively. The cells are sequenced using the inDrop protocol. The filtered BaronMouse dataset has expression data of 1,886 cells and 14,861 genes from 13 cell populations. In the BaronHuman dataset, there are expression data of 8,569 cells and 17,499 genes from 14 cell populations after filtering. The Muraro [27], Segerstolpe [3], and Xin [28] datasets are all from the human pancreas. The Muraro dataset is available on GEO with access number GSE85241 and sequenced by the CEL-Seq2 protocol. After filtering the dataset, there are expression data of 2,122 cells and 18,915 genes, where the cells are annotated to 9 classes. The Segerstolpe dataset includes expression data of 2,133 cells and 22,757 genes, sequenced by the SMART-Seq2 protocol, from 13 cell populations. The last human pancreas dataset is the Xin dataset which is sequenced by the SMARTer protocol. There are 4 types of 1,449 cells in the dataset. For these datasets, we used the cell labels given by the authors as ground truth. Note that the marker genes and computational methods are used to assign these labels.

#### Peripheral blood mononuclear cell

Human peripheral blood mononuclear cells contain heterogeneous cell populations and play an important role in investigating immunology and infectious disease. We used two datasets [29], Zhengsorted and Zheng68K, which were sequenced by the 10x Chromium protocol [30]. In the Zhengsorted dataset, the authors extracted 10 cell populations using antibody based bead enrichment and validated the ten cell types using the FACS sorting. The purified 10 populations were used to generate a large set of single cells individually. Similar to the work in [25], each cell population has 2,000 cells and were mixed together to have an even combined dataset, therefore, there are 20,000 cells of 10 cell classes. The Zheng68K dataset is a large-scale scRNAseq dataset that includes expression of 65,943 cells and 20,387 genes. The cells were labeled by [29] as 11 cell populations. We used the given annotations as ground truth.

### Data preprocessing

All the datasets were filtered by cells and genes. First, we removed the unlabeled cells, and the cells that were labelled as debris and doublets. Also, we removed the genes that have zero expression values across all the cells. Then, we transformed the gene expression values into the log scale and normalized each dataset by min-max scaling. After calculating the variances of the genes across all the cells, we sorted the variances in descending order and chose the top 1,000 genes as the input of the classifiers. For the graph-based method, we constructed the gene adjacency network from the selected genes.

### Complexity of the Datasets

We used the density plot to describe the complexity of the datasets. We calculated the mean expression value of each gene across the cells from the same cell population, as the centroid. Then, we generated a matrix with the dimension of the number of genes by the number of the cell populations for each dataset. The distances between the cell populations were computed using this matrix. Figure 2 shows the density of distance between the centroids of the cell populations. The high peaks of the datasets show that the cell populations are similar to each other and are not easy to separate. We can observe that the Zhengsorted and Zheng68K datasets are the most complex datasets.

**Figure 2:**
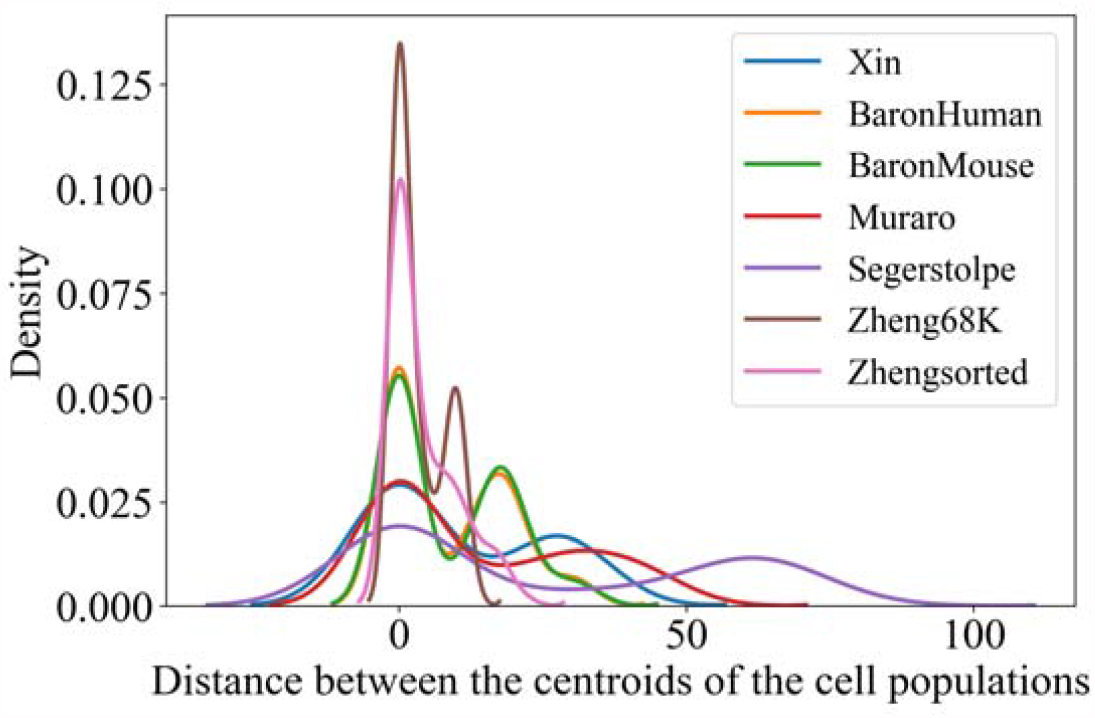
Density plot of the distance between the centroids of cell populations to show the complexity of the datasets.

## Results and discussion

For performance comparison, we used seven tools developed for scRNAseq data classification: scPred [7], scID [8], CaSTLe [9], singleR [12], scmapcluster [13], scmapcell [13], and ACTINN [16]. We also used conventional supervised classifiers including RF, linear SVM (SVM-linear), radial basis function kernel-based SVM (SVM-rbf) and KNN. We used the built-in function in scikit-learn package [31] in Python to implement the traditional classifiers.

To investigate the generalizability of the methods, we used the within-dataset classification and cross-dataset classification to evaluate the performance of the methods. In our within-dataset classification, we randomly split each dataset for training and testing the models. We used the same training and testing datasets for all the methods. In our cross-dataset classification study, first, we combined the four human pancreas datasets (Xin, BaronHuman, Muraro, and Segerstolpe) and then, used three of them as the training dataset and the remaining one as the testing dataset.

### Metrics to evaluate the performance of classification

We used accuracy, F1 score, precision, and recall metrics to evaluate the performance of the classifiers. They are defined as: precision = TP/(TP+FP), recall = TP/(TP+FN), accuracy = (TP+TN)/(TP+TN+FP+FN), F1 = (2 × precision × recall)/(precision + recall), where TP, FP, TN and FN denote true positive, false positive, true negative, and false negative, respectively. We used the accuracy and the median of F1 scores across all the classes to evaluate the overall performance of a classifier. We also showed the median precisions and the median recalls in the results section.

### Performance of within-dataset classification

We evaluated the performance of the classification within each dataset individually. Figure 3 shows the performance of the scRNAseq data classifier tools and the conventional classifiers on the Zhengsorted dataset in comparison to our proposed classifier, sigGCN, in terms of accuracy, F1, precision, and recall. Note that the Zhengsorted dataset is the only dataset with validated ground truth cell types. For other datasets, the labels are obtained by employing a clustering method on the marker genes. The accuracy (0.922), median F1 (0.965), precision (0.970), and recall (0.972) of our model are improved compared to not only the traditional machine learning methods but also the methods that are developed specifically for cell classification. Our model performs better than the fully connected neural network based method, i.e. ACTINN (accuracy=0.845, F1 score=0.892, precision = 0.913, recall = 0.886).

**Figure 3:**
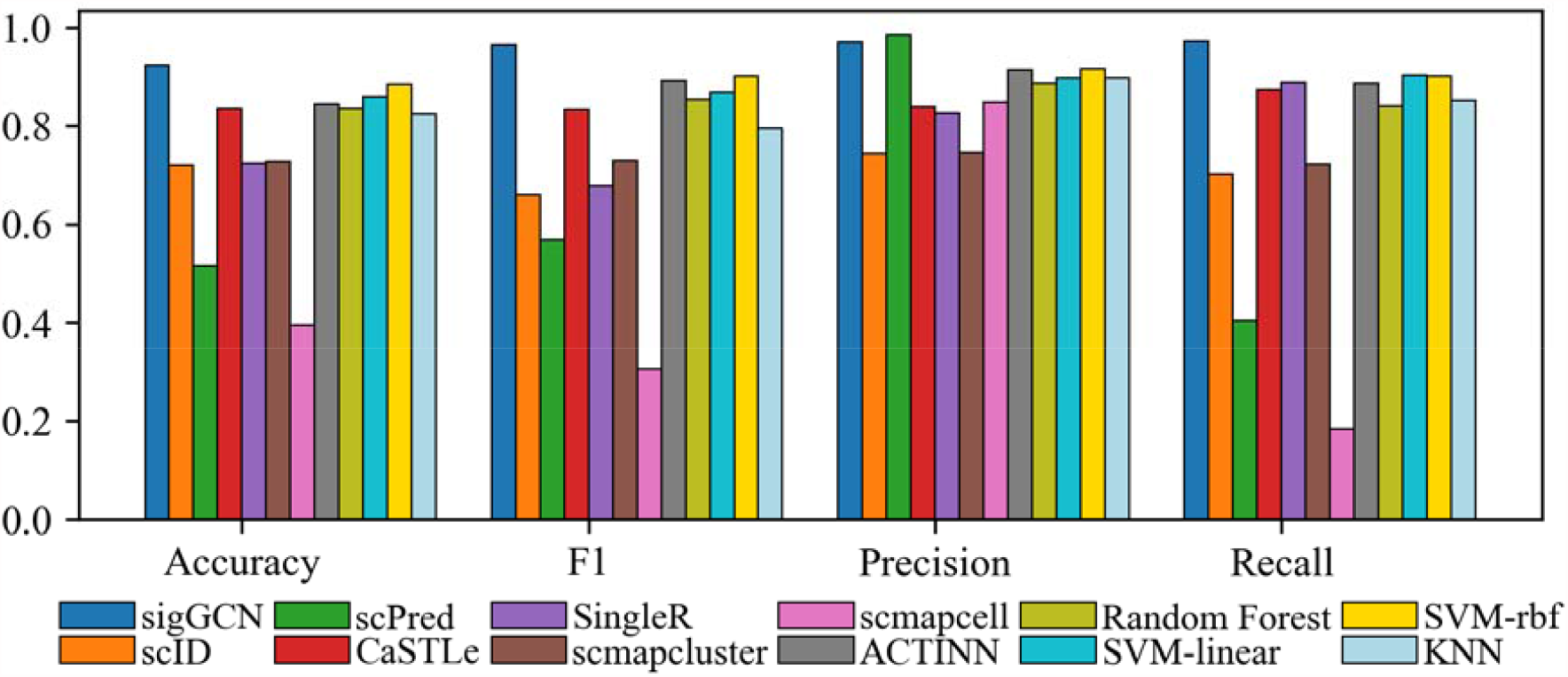
Bar plots of the four metrics to show the performance of the scRNAseq data classifier tools and conventional classifiers on Zhengsorted dataset.

The performances of all the methods on the Zheng68K dataset with 11 classes are shown in Additional file 1: Figure S1. As can be seen, sigGCN performs better than the other methods with an accuracy of 0.752, median F1 of 0.776, precision of 0.735, and recall of 0.780. Notice that all the methods do not perform well using the Zheng68K dataset because it is the most complex dataset as shown in Figure 2.

The results show that our deep learning based method performs better than the conventional machine learning classification methods (RF, SVM, and KNN) using the Zhengsorted and Zheng68K dataset. One reason is that the conventional methods are not scalable for large datasets. The Zhengsorted dataset with 20,000 cells is a large scale scRNAseq dataset which benefits the deep learning based methods.

The performances of all the methods using the other five datasets in terms of accuracy, F1, precision, and recall are shown in the Additional file 1: Figures S2-S6. Table 1 and Table 2 show the accuracy and median F1 score of all the methods using the seven datasets, respectively. The results show that our model (sigGCN) outperforms the scRNAseq classifier tools and the conventional classifier methods using most of the datasets.

**Table 1.** Accuracy of the eight scRNAseq data classifier tools and the four conventional classifiers on the seven datasets (*N*=1000)

| Methods | Zhengsorted | Zheng68K | BaronHuman | Muraro | Segerstolpe | BaronMouse | Xin |
| --- | --- | --- | --- | --- | --- | --- | --- |
| sigGCN | **0.922** | **0.752** | **0.979** | **0.991** | **0.977** | 0.974 | 0.993 |
| scID | 0.721 | 0.484 | 0.46 | 0.577 | 0.285 | 0.286 | 0.986 |
| scPred | 0.515 | 0.140 | 0.86 | 0.915 | 0.827 | 0.862 | 0.91 |
| CasTLe | 0.836 | 0.736 | 0.971 | 0.972 | 0.953 | 0.91 | 0.993 |
| SingleR | 0.723 | 0.388 | 0.951 | 0.977 | 0.953 | 0.868 | **1** |
| scmapcluster | 0.395 | 0.409 | 0.946 | 0.962 | 0.949 | 0.905 | 0.931 |
| scmapcell | 0.727 | 0.246 | 0.895 | 0.972 | 0.949 | 0.778 | 0.952 |
| ACTINN | 0.845 | 0.737 | 0.977 | 0.991 | 0.958 | **0.984** | **1** |
| RF | 0.835 | 0.69 | 0.968 | 0.981 | 0.967 | 0.968 | 0.993 |
| SVM-linear | 0.859 | 0.652 | 0.824 | 0.981 | 0.383 | 0.704 | **1** |
| SVM-rbf | 0.884 | 0.677 | 0.93 | 0.981 | 0.841 | 0.794 | **1** |
| KNN | 0.824 | 0.594 | 0.953 | 0.991 | 0.935 | 0.926 | **1** |

**Table 2.** Median F1 of the eight scRNAseq data classifier tools and the four conventional classifiers on the seven datasets (*N*=1000)

| Methods | Zhengsorted | Zheng68K | BaronHuman | Muraro | Segerstolpe | BaronMouse | Xin |
| --- | --- | --- | --- | --- | --- | --- | --- |
| sigGCN | **0.965** | **0.776** | 0.977 | **1** | **1** | 0.969 | 0.995 |
| scID | 0.66 | 0.535 | 0.22 | 0.578 | 0 | 0 | **1** |
| scPred | 0.568 | 0.105 | 0.833 | 0.932 | 0.8 | **0.97** | 0.784 |
| CasTLe | 0.834 | 0.667 | 0.956 | 0.967 | 0.965 | 0.848 | 0.997 |
| SingleR | 0.678 | 0.335 | 0.946 | 0.984 | **1** | 0.898 | **1** |
| scmapcluster | 0.729 | 0.357 | 0.9 | 0.997 | 0.965 | 0.88 | 0.991 |
| scmapcell | 0.305 | 0.198 | 0.95 | 0.993 | 0.977 | 0.942 | 0.708 |
| ACTINN | 0.892 | 0.753 | **1** | 0.97 | **1** | **1** | 0.997 |
| RF | 0.853 | 0.646 | 0.956 | 0.987 | 0.993 | 0.984 | 0.997 |
| SVM-linear | 0.868 | 0.663 | 0.362 | **1** | 0.059 | 0.238 | **1** |
| SVM-rbf | 0.9 | 0.671 | 0.906 | **1** | 0.772 | 0.695 | **1** |
| KNN | 0.795 | 0.613 | 0.928 | **1** | 0.961 | 0.889 | **1** |

Almost all the methods perform well on the Xin, Segerstolpe, and Muraro datasets and there are not significant differences between the performance of the methods. It can be due to well-separated cell population in these datasets. As can be seen in Figure 2 the distances between clusters are higher in these datasets compared to the rest. Interestingly, for more complex datasets, Zhengsorted, Zheng68K, in which the Euclidian distances between clusters are lower, our proposed method performs significantly better as can be seen in Table 1.

Using the BaronHuman dataset, our model shows the best performance against other methods with an accuracy of 0.979 and median F1 of 0.977. The RF and KNN have good accuracy but could not hold the high median precision and median recall values. This is because the accuracy is calculated based on the global TPs and TNs, but the medians of all the paired scores indicate that some classes with a small number of members were not classified well using the conventional methods. Our model reduces FN predictions and increases TP predictions, which notably improves the median recall. In addition, the FPs of our model are also reduced that substantially enhances the precision. Therefore, the median F1, median precision and recall are higher compared to those of the other methods using the BaronHuman dataset.

The performance of our model using the BaronMouse dataset is comparable to that of the ACTINN and better than those of the other methods. Overall, our proposed model shows the best or near the best performance using all the datasets, especially for complex datasets that have large cell numbers and more classes. The performance of sigGCN is improved significantly in comparison to the pure neural network which indicates that integrating the prior knowledge of gene interactions can learn a better representation of data. The feature representation learned by convolution through the gene interaction structure is not redundant with that learned through the expression values. The parallel structure enriches extracting effective features during the training.

We also evaluated the performance of the proposed method when constructing the gene graph using i) the gene cosine similarity, and ii) the gene interaction network without singleton genes. Results show that employing the gene interaction network (including singleton genes) provides the best performance (as can be seen from Additional file 1: Tables S1 – S4). To evaluate the performance of the proposed parallel network compared to employing only a GCN, we compared the performance of sigGCN with that of a pure GCN. Results in Additional file 1: Table S5 show that the parallel network outperforms the pure GCN model, which indicates that the parallel network (NN and GCN) can capture global and local features.

We need to mention that scPred, scID, scmapcluster, and scmapcell provide the function of rejection which means they predict a cell class as “unassigned”. We computed the accuracy and F1 scores based on the results of including these unassigned cells in our comparison (Table 1 and Table2). Since our method outputs the probability of cell class assignments, we also provide an additional function to predict a cell class as “unassigned” by setting a threshold of prediction. We set the threshold as 0.65 which means if a cell does not have a probability of prediction larger than 0.65, then the cell will be predicted as “unassigned”. We computed accuracy and F1 scores when not including the unassigned cells for these four methods and our method with the “unassigned” function (shown in Additional file 2 Table). Results show that our method has the smallest unassigned rate and the best or near the best accuracy and F1.

### Confusion matrix and ROC analysis

Since the Zhengsorted dataset is the one that has the ground truth, we provided more details on the performance of the methods using this dataset in this section. The confusion matrix of the ten classes using our proposed method, sigGCN, is shown in Figure 4. The confusion matrices of the other scRNAseq classifiers are shown in Additional file 1: Figures S7-S8, and those of the conventional methods are shown in Additional file 1: Figure S9. As can be seen in Figure 4, using our model the TP rate of class 0, to class 9 are 99.5%, 100%, 99.5%, 69.3%, 80.9%, 84.6%, 97.1%, 99.0%, 97.4%, 92.0%, respectively. Class 3, class 4, and class 5 have confusion with each other because CD4+ cells have high heterogeneity.

**Figure 4:**
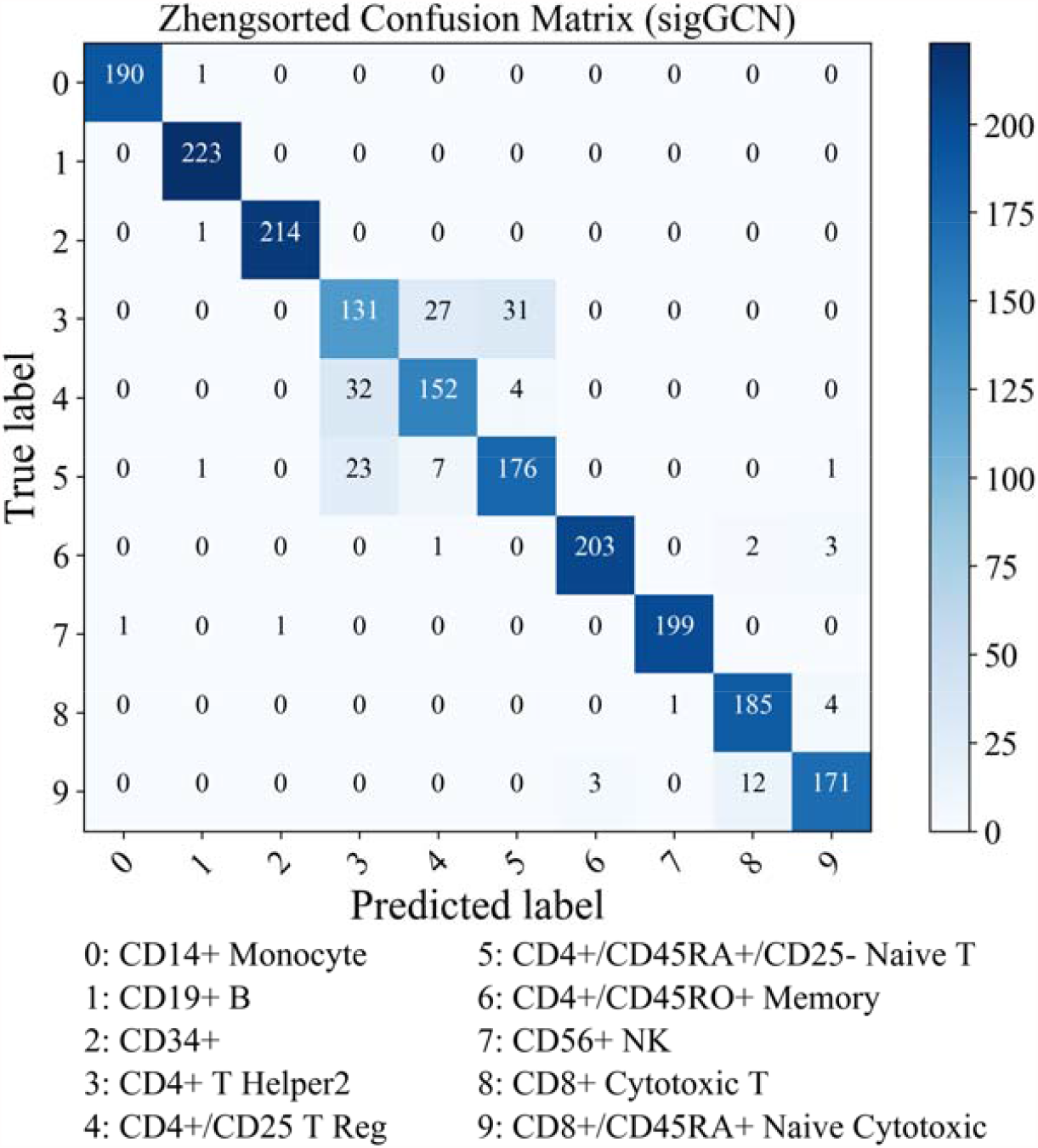
Confusion matrix of the class predictions on the Zhengsorted dataset using sigGCN.

To further investigate the performance of the methods, we executed the ROC analysis for each class using the Zhengsorted dataset. Figure 5(a) shows the ROCs for each class using our proposed method. We also compared our model with the other methods in terms of AUC and the ROC analysis of each class. The ROC analysis for class 4 (CD4+/CD25 T Reg) using all the methods is shown in Figure 5(b). We only showed the performance of six methods (two scRNAseq classifiers and four conventional classifiers) which provides the output classification probability. Our model shows the best AUC of 0.9872 in the ROC analysis of class 4. To test the significance of the difference between the AUCs under the curves, we used the McNeil & Hanley’s test [32] and online service on http://vassarstats.net/roc_comp.html. The p-values of the tests between our method sigGCN and each method are shown in Figure 5(b). The ROC analysis of class 3, class 5, class 6, and class 8 are also shown in Additional file 1: Figure S10.

**Figure 5:**
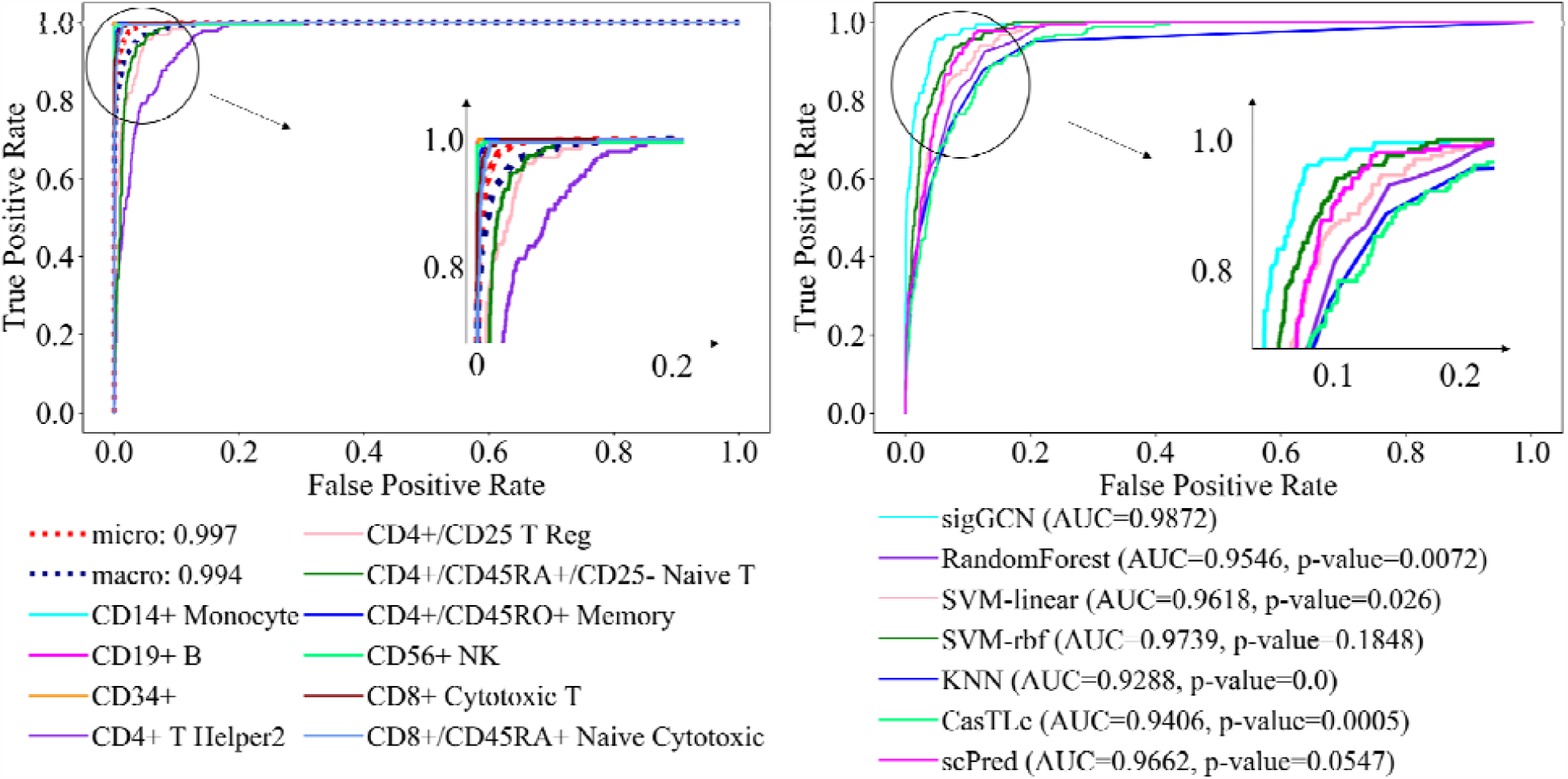
(a) Average AUC and ROC curves of sigGCN using the ten Zhengsorted cell types. (b) ROC curves of class 4 (CD4+/CD25 T Reg) and the p-values calculated by the McNeil & Hanley’s test that show the significance of difference between the areas under the ROC curve of sigGCN and that of each method.

### Performance of cross-dataset classification

In order to examine the generalization of the proposed model, we evaluated the performance of cross-dataset classification, which is a more realistic scenario. Since the Xin, BaronHuman, Muraro, and Segerstolpe datasets are all from the human pancreas, we used these four datasets for the validation. The common cell types among these four datasets are alpha, beta, delta, and gamma, so we extracted the four cell types from each dataset for combination. Before combining the datasets, we preprocessed the data using the log-transformation and normalized each dataset by min-max scaling to make the four datasets in the same level. We run four experiments, and in each experiment, we used three of the four datasets as the training dataset and the remaining one as the testing dataset. Additional file 1: Figures S11-S14 show the performance of the methods in terms of accuracy, median F1, median precision, and median recall score. The results of the accuracy are shown in Table 3 and the median F1 scores are shown in Table 4, respectively. We included the unassigned cells when computing the accuracy and F1 scores for the scPred, scID, scmapcluster, and scmapcell. We also provided the results of not including the unassigned cells in Additional file 3.

**Table 3.** Accuracy of the eight scRNAseq data classifier and the four conventional classifiers tools on the four experiments (*N*=1000)

| Training Dataset | BaronHuman+<br>Muraro+ Segerstolpe | Xin+ Muraro+<br>Segerstolpe | Xin+ BaronHuman+<br>Segerstolpe | Xin+<br>BaronHuman+<br>Muraro |
| --- | --- | --- | --- | --- |
| Testing Dataset | Xin | BaronHuman | Muraro | Segerstolpe |
| sigGCN | **0.997** | **0.987** | 0.974 | 0.993 |
| scID | 0.989 | 0.747 | 0.97 | 0.979 |
| scPred | 0.945 | 0.467 | 0.92 | 0.814 |
| CasTLe | 0.99 | 0.944 | **0.977** | 0.992 |
| SingleR | 0.995 | 0.984 | **0.977** | **0.996** |
| scmapcluster | 0.196 | 0.003 | 0.051 | 0.568 |
| scmapcell | 0.756 | 0.421 | 0.64 | 0.367 |
| ACTINN | 0.993 | 0.984 | 0.974 | 0.992 |
| RF | 0.982 | 0.941 | 0.947 | 0.938 |
| SVM-linear | 0.994 | 0.979 | 0.972 | 0.992 |
| SVM-rbf | 0.986 | 0.983 | 0.973 | 0.97 |
| KNN | 0.934 | 0.864 | 0.788 | 0.817 |

**Table 4.**
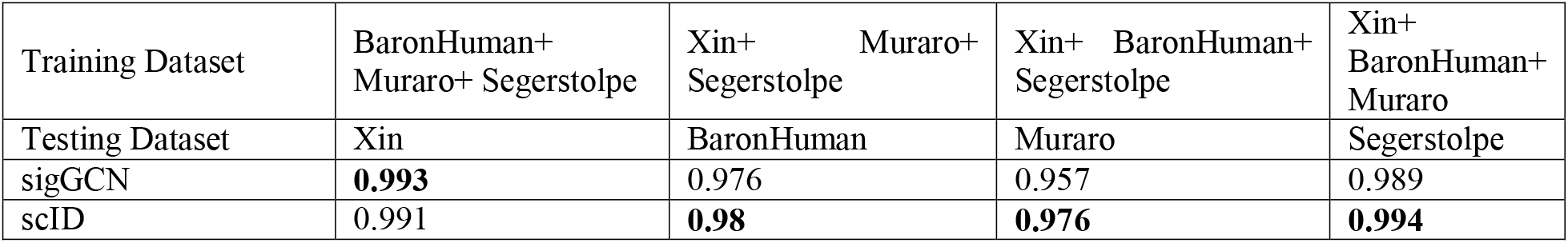
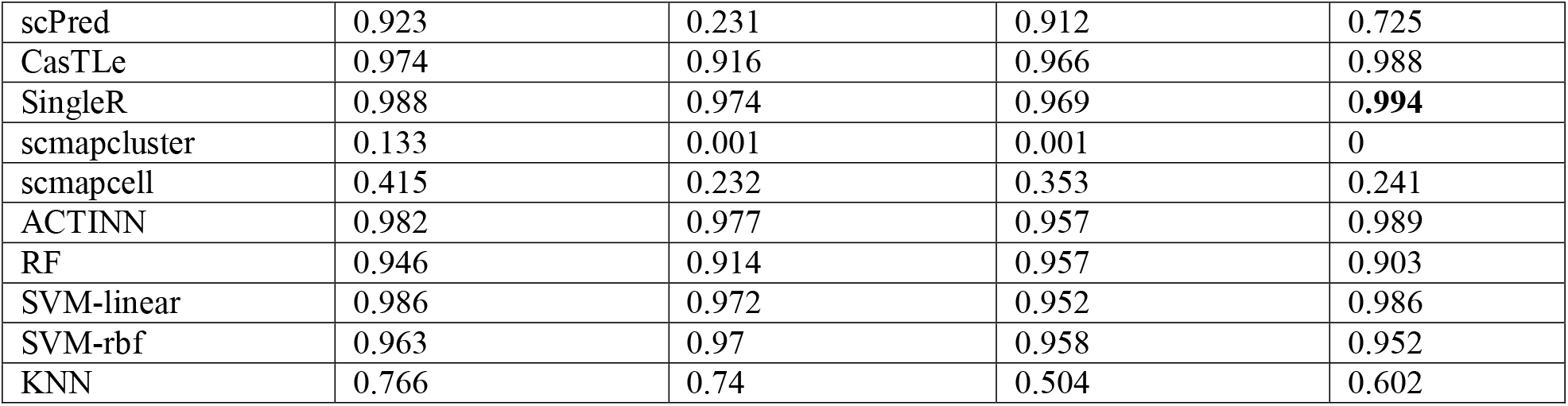
Median F1 of the eight scRNAseq data classifier and the four conventional classifiers tools on the four experiments (*N*=1000)

All the methods except the scmapcluster, scmapcell, and KNN perform nearly perfect in terms of accuracy when training on the BaronHuman, Muraro, and Segerstolpe and testing on the Xin dataset. Providing a near 100% accuracy, F1, precision, and recall, our model shows the best performance compared to the other methods. Our model ranks the third and the second in terms of accuracy when testing on the Muraro and Segerstolpe datasets, respectively, with a slight difference of 0.003. Overall, our model achieves the best or near the best performance in terms of accuracy and median F1 in the cross-dataset classification evaluation.

### Performance of runtimes

We compared the runtimes of our method and the other scRNAseq classifiers. The runtimes were computed by using an iMac with a 4.2GHz Intel Core i7 CPU and up to 32 gigabytes of memory. The average runtime of each method using the Zhengsorted dataset is shown in Figure 6. The runtimes of all the methods using the other six datasets are shown in Additional file 1: Figure S15. The average runtimes of sigGCN are 404.3s, 1237.8s, 164.0s, 41.9s, 31.3s, 35.5s, 19.7s using Zhengsorted, Zheng68K, Baron Human, Muraro, Segerstolpe, Baron Mouse, and Xin datasets, respectively. For the five pancreas dataset that have a small number of cells, all the methods run less than 10 mins. For the large scale datasets, Zhengsorted with 20,000 cells and Zheng68K with more than 60,000 cells, the average runtimes of our method are less than or comparable to those of scID, CasTLe, singleR, and ACTINN. The runtimes of scmapcluster and scmapcell are always low, even on the Zheng68K dataset, because scmapcluster assigns the labels only by computing the distance between the testing cells and a small group of virtual cells.

**Figure 6:**
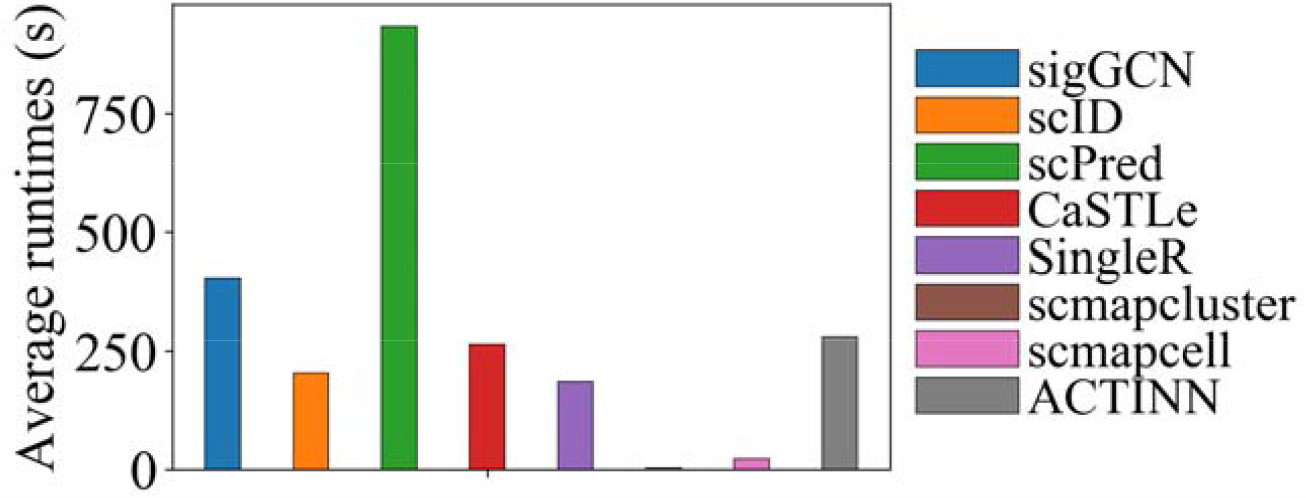
Average runtimes of the methods using Zhengsorted dataset.

### Evaluation of sensitivity to the number of genes

We evaluated the sensitivity of the methods to the number of input genes in this section. We used the same subset of genes which was selected based on the gene expression variance across the Zhengsorted dataset as the input of all the scRNAseq classifiers including sigGCN. We selected top 300, 500, 750, 1000, 1250, 1500, and 5000 highest variant genes. The accuracy and median F1 of using a different number of genes using the Zhengsorted dataset are shown in Figure 7. As can be seen, our method, sigGCN, has a consistent performance in terms of accuracy and median F1 when using different sizes of the input features (high variant genes). The model performance for using a large size of genes (> 5,000) does not show a significant improvement compared with that of using a small size of genes (≤ 1,000). Considering the time complexity, we selected the top 1,000 variable genes as the default input in our method.

**Figure 7:**
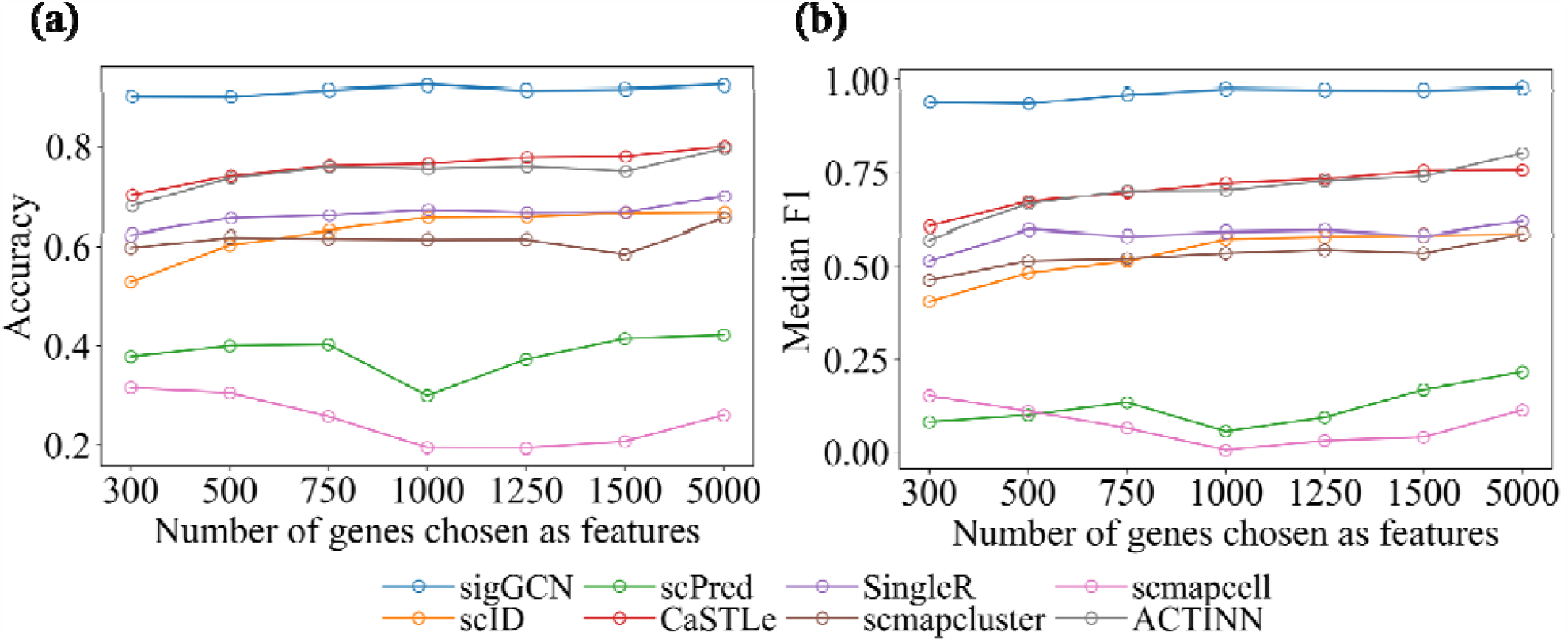
Results of choosing different numbers of genes as features of all the scRNAseq classifiers using Zhengsorted dataset.

### Evaluation of scalability to the number of cells

To investigate the model scalability to different size of the dataset, we down-sampled the Zhengsorted dataset into five subsets with the sampling rate of 0.05, 0.1, 0.15, 0.25, 0.5. The ten cell classes of the Zhengsorted dataset were down-sampled in a stratified way to guarantee that each cell class has the same percentage. Therefore, the five subsets of Zhengsorted dataset have 1,000 cells, 2,000 cells, 3,000 cells, 5,000 cells, and 10,000 cells. We divided each down-sampled subset to 80% as the training dataset, 10% as the validation dataset, and 10% as the testing dataset. The performance in terms of accuracy and median F1 of the scRNAseq classifiers is shown in Figure 8. In general, the accuracy and median F1 scores of all the methods increase as the number of cells increases. Our method can hold consistent median F1 on all the down-sampled subsets.

**Figure 8:**
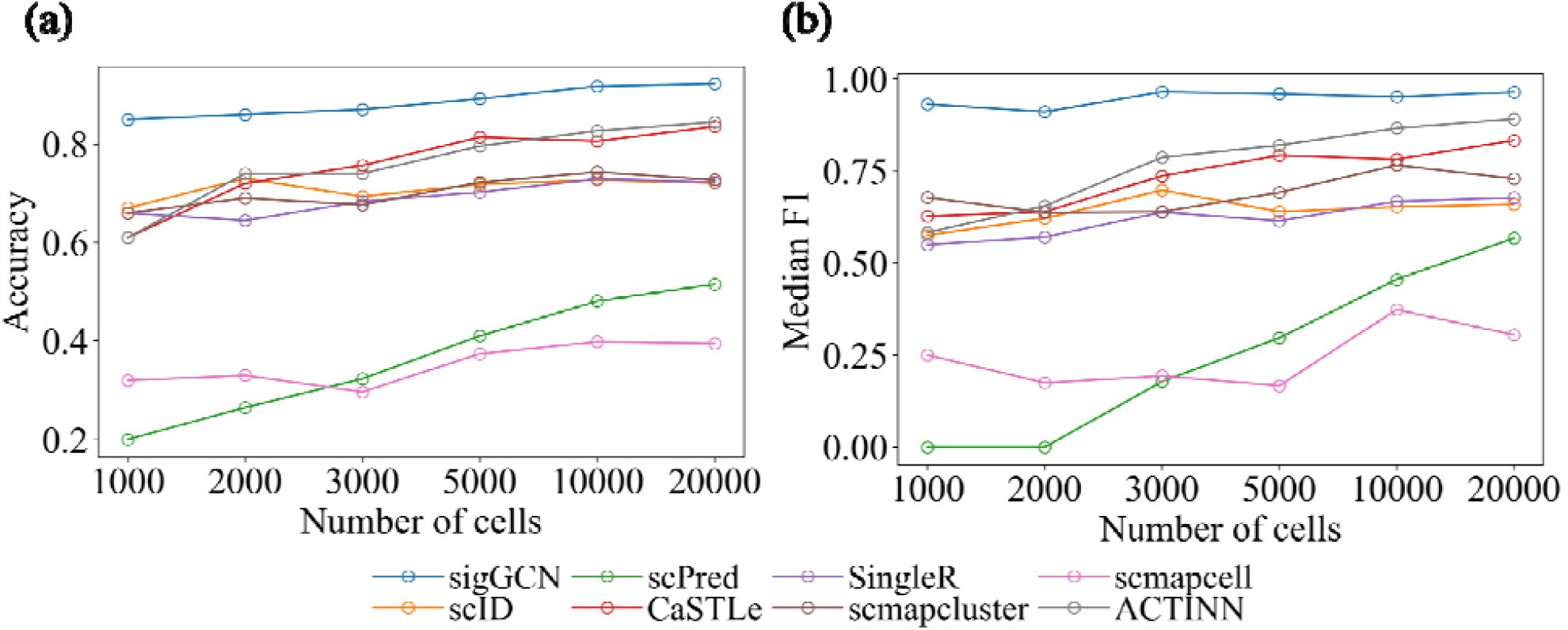
Results of all the scRNAseq classifiers using different numbers of cells from the Zhengsorted dataset.

## Discussion and Conclusion

In this study, we propose a novel method based on the deep learning methodologies for cell classification in single cell studies when a large number of cells are used and data are heterogeneous with many classes. We integrated the prior knowledge, in the form of gene-gene interaction network, into the classification procedure. We developed an end-to-end trained multimodal deep learning model which includes a GCN and an NN. The GCN model exploits the gene network and gene expression values to extract the integrated features regarding the interconnection between genes, while the NN model is utilized to enrich the features extracted by the GCN model.

We compared the performance of our proposed method with those of seven other classification methods and four traditional machine learning classification methods using seven real single cell RNAseq datasets. The standard classification metrics, including accuracy, median F1, median precision, and median recall scores were utilized to evaluate the performance of the methods on all the 7 datasets. We further evaluated the classification performance on a large and highly heterogeneous dataset, the Zhengsorted dataset, by comparing the confusion matrix and the ROC plot. Results show that our proposed method, sigGCN, outperforms both the current scRNAseq classification tools and the conventional machine learning methods.

The main contribution of this work is integrating the prior knowledge in the form of the interaction between genes into the classification of cell types. By aggregating the features from neighboring nodes in the graph, the new features are learned in a nonlinear manner. The feature representation of the data using a GCN is not redundant with that learned by using only an NN due to convolution through the gene interaction structure. Therefore, we proposed to utilize a parallel structure of a GCN and an NN to integrate the gene interaction structure and gene expression values.

In conclusion, the proposed multimodal deep learning model which integrates gene expression data with the biological network to classifying cells shows a powerful performance. In the future work, we will introduce the attention mechanism to enhance the weights of important genes.

## List of abbreviations

scRNAseq: Single Cell RNA Sequencing
GCN: Graph convolutional network
NN: Neural network
ROC: Receiver Operating Characteristic
TP: True Positive
FP: False Positive
TN: True Negative
FN: False Negative
AUC: Area Under Curve

## Declarations

### Ethics approval and consent to participate

No ethics approval was required for the study.

## Consent for publication

Not applicable.

## Availability of data and materials

The single cell RNAseq data are downloaded from https://doi.org/10.5281/zenodo.3357167 [25]. The code is available at https://github.com/NabaviLab/sigGCN.

## Authors’ contributions

SN and TW designed the study. TW and JB designed the models. TW collected the data and implemented the analysis. TW, SN, and JB wrote the manuscript. All authors read and confirmed the manuscript.

## Acknowledgments

Not applicable.

## Additional Files

Additional file 1: Supplementary materials (Supplementary Tables S1-S4, Supplementary Figures S1-S15).

Additional file 2: Additional_file_2_results_within-dataset_excluding_unassigned_cells, results of the classification accuracy, F1 scores, and unassigned rate of not including the unassigned cells for five methods (scPred, scID, scmapcluster, scmapcell, and sigGCN) using within-dataset experiments.

Additional file 3: Additional_file_2_results_cross-dataset_excluding_unassigned_cells, results of the classification accuracy, F1 scores, and unassigned rate of not including the unassigned cells for five methods (scPred, scID, scmapcluster, scmapcell, and sigGCN) using cross-dataset experiments.

## References

1. Villani A-C, Satija R, Reynolds G, Sarkizova S, Shekhar K, Fletcher J, et al. Single-cell RNA-seq reveals new types of human blood dendritic cells, monocytes, and progenitors. Science. 2017;356. doi:10.1126/science.aah4573.

2. Grün D, Lyubimova A, Kester L, Wiebrands K, Basak O, Sasaki N, et al. Single-cell messenger RNA sequencing reveals rare intestinal cell types. Nature. 2015;525:251–5.

3. Segerstolpe Å, Palasantza A, Eliasson P, Andersson E-M, Andréasson A-C, Sun X, et al. Single-Cell Transcriptome Profiling of Human Pancreatic Islets in Health and Type 2 Diabetes. Cell Metab. 2016;24:593–607.

4. Fincher CT, Wurtzel O, Hoog T de, Kravarik KM, Reddien PW. Cell type transcriptome atlas for the planarian Schmidtea mediterranea. Science. 2018;360. doi:10.1126/science.aaq1736.

5. Plass M, Solana J, Wolf FA, Ayoub S, Misios A, Glažar P, et al. Cell type atlas and lineage tree of a whole complex animal by single-cell transcriptomics. Science. 2018;360. doi:10.1126/science.aaq1723.

6. Zhao X, Wu S, Fang N, Sun X, Fan J. Evaluation of single-cell classifiers for single-cell RNA sequencing data sets. Brief Bioinform. 2019;:bbz096.

7. Alquicira-Hernandez J, Sathe A, Ji HP, Nguyen Q, Powell JE. scPred: accurate supervised method for cell-type classification from single-cell RNA-seq data. Genome Biol. 2019;20:264.

8. Boufea K, Seth S, Batada NN. scID: Identification of equivalent transcriptional cell populations across single cell RNA-seq data using discriminant analysis. bioRxiv. 2019;:470203.

9. Lieberman Y, Rokach L, Shay T. CaSTLe – Classification of single cells by transfer learning: Harnessing the power of publicly available single cell RNA sequencing experiments to annotate new experiments. PLOS ONE. 2018;13:e0205499.

10. Tan Y, Cahan P. SingleCellNet: a computational tool to classify single cell RNA-Seq data across platforms and across species. preprint. Bioinformatics; 2018. doi:10.1101/508085.

11. Chen T, Guestrin C. XGBoost: A Scalable Tree Boosting System. Proc 22nd ACM SIGKDD Int Conf Knowl Discov Data Min. 2016;:785–94.

12. Aran D, Looney AP, Liu L, Wu E, Fong V, Hsu A, et al. Reference-based analysis of lung single-cell sequencing reveals a transitional profibrotic macrophage. Nat Immunol. 2019;20:163–72.

13. Kiselev VY, Yiu A, Hemberg M. scmap: projection of single-cell RNA-seq data across data sets. Nat Methods. 2018;15:359–62.

14. Wagner F, Yanai I. Moana: A robust and scalable cell type classification framework for single-cell RNA-Seq data. preprint. Bioinformatics; 2018. doi:10.1101/456129.

15. Zhang Z, Danni Luo MS, Zhong X, Choi JH, Ma Y, Mahrt E, et al. SCINA: Semi-Supervised Analysis of Single Cells in silico. preprint. Bioinformatics; 2019. doi:10.1101/559872.

16. Ma F, Pellegrini M. ACTINN: automated identification of cell types in single cell RNA sequencing. Bioinforma Oxf Engl. 2020;36:533–8.

17. Wu Z, Pan S, Chen F, Long G, Zhang C, Yu PS. A Comprehensive Survey on Graph Neural Networks. IEEE Trans Neural Netw Learn Syst. 2020;:1–21.

18. Defferrard M, Bresson X, Vandergheynst P. Convolutional neural networks on graphs with fast localized spectral filtering. In: Proceedings of the 30th International Conference on Neural Information Processing Systems. Barcelona, Spain: Curran Associates Inc.; 2016. p. 3844–52.

19. Fout A, Byrd J, Shariat B, Ben-Hur A. Protein Interface Prediction using Graph Convolutional Networks. In: Guyon I, Luxburg UV, Bengio S, Wallach H, Fergus R, Vishwanathan S, et al., editors. Advances in Neural Information Processing Systems 30. Curran Associates, Inc.; 2017. p. 6530–9. http://papers.nips.cc/paper/7231-protein-interface-prediction-using-graph-convolutional-networks.pdf. Accessed 1 Jun 2020.

20. Sun M, Zhao S, Gilvary C, Elemento O, Zhou J, Wang F. Graph convolutional networks for computational drug development and discovery. Brief Bioinform. 2020;21:919–35.

21. Szklarczyk D, Gable AL, Lyon D, Junge A, Wyder S, Huerta-Cepas J, et al. STRING 11: protein–protein association networks with increased coverage, supporting functional discovery in genome-wide experimental datasets. Nucleic Acids Res. 2019;47:D607–13.

22. Shuman DI, Narang SK, Frossard P, Ortega A, Vandergheynst P. The Emerging Field of Signal Processing on Graphs: Extending High-Dimensional Data Analysis to Networks and Other Irregular Domains. IEEE Signal Process Mag. 2013;30:83–98.

23. Hammond DK, Vandergheynst P, Gribonval R. Wavelets on Graphs via Spectral Graph Theory. ArXiv09123848 Cs Math. 2009. http://arxiv.org/abs/0912.3848. Accessed 18 May 2020.

24. Ruder S. An overview of gradient descent optimization algorithms. ArXiv160904747 Cs. 2017. http://arxiv.org/abs/1609.04747. Accessed 5 Dec 2020.

25. Abdelaal T, Michielsen L, Cats D, Hoogduin D, Mei H, Reinders MJT, et al. A comparison of automatic cell identification methods for single-cell RNA sequencing data. Genome Biol. 2019;20:194.

26. Baron M, Veres A, Wolock SL, Faust AL, Gaujoux R, Vetere A, et al. A Single-Cell Transcriptomic Map of the Human and Mouse Pancreas Reveals Inter-and Intra-cell Population Structure. Cell Syst. 2016;3:346-360.e4.

27. Muraro MJ, Dharmadhikari G, Grün D, Groen N, Dielen T, Jansen E, et al. A Single-Cell Transcriptome Atlas of the Human Pancreas. Cell Syst. 2016;3:385-394.e3.

28. Xin Y, Kim J, Okamoto H, Ni M, Wei Y, Adler C, et al. RNA Sequencing of Single Human Islet Cells Reveals Type 2 Diabetes Genes. Cell Metab. 2016;24:608–15.

29. Zheng GXY, Terry JM, Belgrader P, Ryvkin P, Bent ZW, Wilson R, et al. Massively parallel digital transcriptional profiling of single cells. Nat Commun. 2017;8:14049.

30. 10x Genomics: Resolving Biology to Advance Human Health. https://www.10xgenomics.com/. Accessed 10 Jan 2020.

31. Pedregosa F, Varoquaux G, Gramfort A, Michel V, Thirion B, Grisel O, et al. Scikit-learn: Machine Learning in Python. Mach Learn PYTHON. :6.

32. Hanley JA, McNeil BJ. The meaning and use of the area under a Receiver Operating Characteristic (ROC) curve. Radiology. 1982;143:29–36.

