## Additional file 1 for "Single-Cell Classification Using Graph Convolutional Networks"

**Supplementary**

**Supplement tables**

**Table S1. Accuracy of using different gene networks on the seven datasets**

| Methods | Zhengsorted | Zheng68K | BaronHuman | Muraro | Segerstolpe | BaronMouse | Xin |
| --- | --- | --- | --- | --- | --- | --- | --- |
| sigGCN | 0.922 | 0.752 | 0.979 | 0.991 | 0.977 | 0.974 | 0.993 |
| sigGCN-exprSimilarity | 0.905 | 0.751 | 0.973 | 0.986 | 0.967 | 0.958 | 0.993 |
| sigGCN-w/oSingleton | 0.904 | 0.752 | 0.977 | 0.986 | 0.972 | 0.963 | 0.993 |

**Table S2. Median F1 of using different gene networks on the seven datasets**

| Methods | Zhengsorted | Zheng68K | BaronHuman | Muraro | Segerstolpe | BaronMouse | Xin |
| --- | --- | --- | --- | --- | --- | --- | --- |
| sigGCN | 0.965 | 0.776 | 0.977 | 1 | 1 | 0.969 | 0.995 |
| sigGCN- exprSimilarity | 0.949 | 0.773 | 0.965 | 1.000 | 0.951 | 0.937 | 0.995 |
| sigGCN- w/oSingleton | 0.941 | 0.783 | 0.963 | 1.000 | 0.963 | 0.964 | 0.995 |

**Table S3. Accuracy of using different gene networks on the four experiments**

| Training Dataset | BaronHuman+ Muraro+ Segerstolpe | Xin+ Muraro+ Segerstolpe | Xin+ BaronHuman+ Segerstolpe | Xin+ BaronHuman+ Muraro |
| --- | --- | --- | --- | --- |
| Testing Dataset | Xin | BaronHuman | Muraro | Segerstolpe |
| sigGCN | 0.997 | 0.987 | 0.974 | 0.993 |
| sigGCN- exprSimilarity | 0.996 | 0.982 | 0.973 | 0.990 |
| sigGCN- w/oSingleton | 0.995 | 0.980 | 0.965 | 0.985 |

**Table S4. Median F1 of using different gene networks on the four experiments**

| Training Dataset | BaronHuman+ Muraro+ Segerstolpe | Xin+ Muraro+ Segerstolpe | Xin+ BaronHuman+ Segerstolpe | Xin+ BaronHuman+ Muraro |
| --- | --- | --- | --- | --- |
| Testing Dataset | Xin | BaronHuman | Muraro | Segerstolpe |
| sigGCN | 0.993 | 0.976 | 0.957 | 0.989 |
| sigGCN- exprSimilarity | 0.993 | 0.965 | 0.956 | 0.980 |
| sigGCN- w/oSingleton | 0.989 | 0.968 | 0.943 | 0.971 |

**Table S5. Accuracy of using the proposed parallel model sigGCN, and pure GCN with two different loss functions on the seven datasets.**

| Methods | Zhengsorted | Zheng68K | BaronHuman | Muraro | Segerstolpe | BaronMouse | Xin |
| --- | --- | --- | --- | --- | --- | --- | --- |
| sigGCN | 0.922 | 0.752 | 0.979 | 0.991 | 0.977 | 0.974 | 0.993 |
| GCNrecon | 0.873 | 0.66 | 0.968 | 0.991 | 0.949 | 0.958 | 0.993 |
| GCNpure | 0.852 | 0.656 | 0.969 | 0.991 | 0.949 | 0.937 | 0.986 |

*GCNrecon is a GCN with cross entropy loss and reconstruction loss; and GCNpure is a GCN with cross entropy loss.

**Supplement figures**

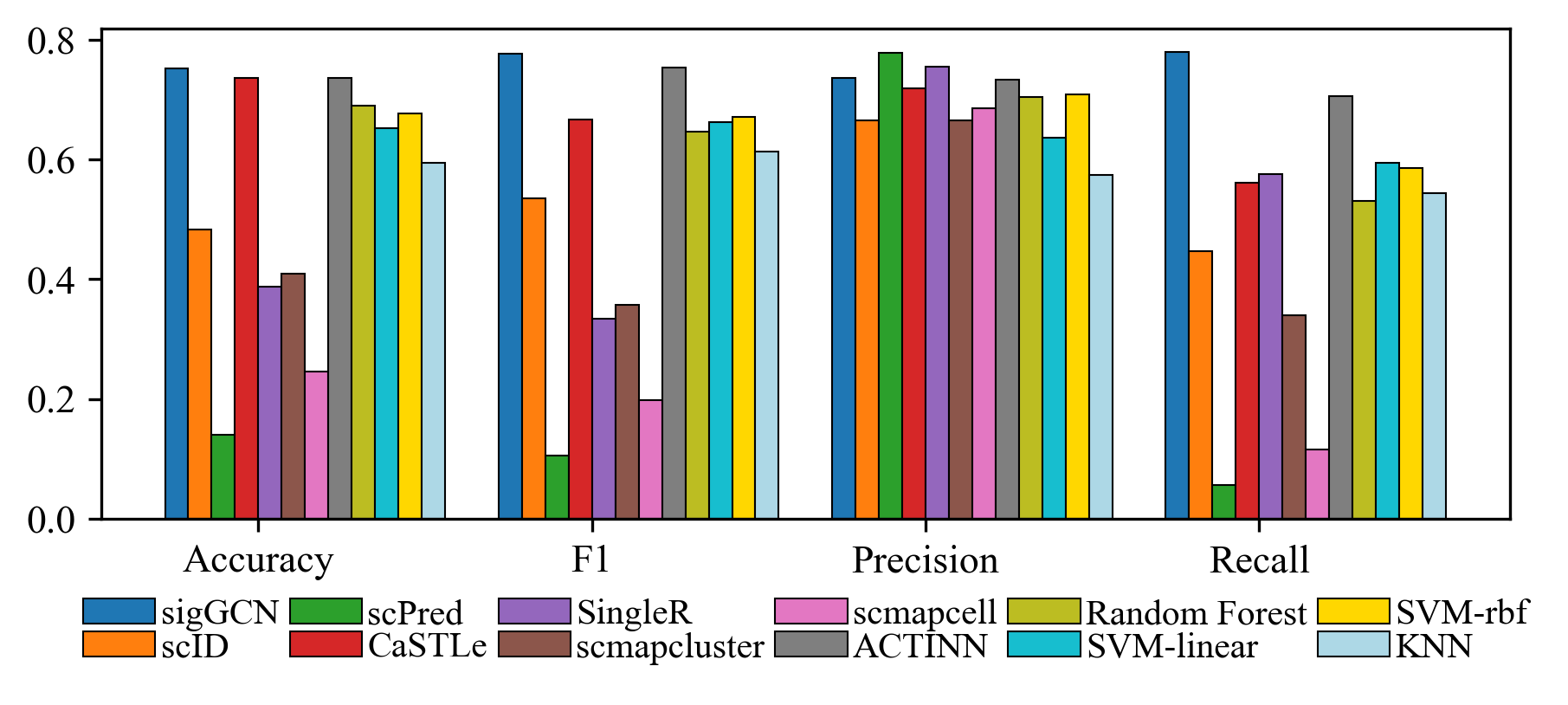

Figure S1: Bar plots of the four metrics to show the performance of scRNAseq data classifier tools and conventional classifiers on Zheng68K dataset.

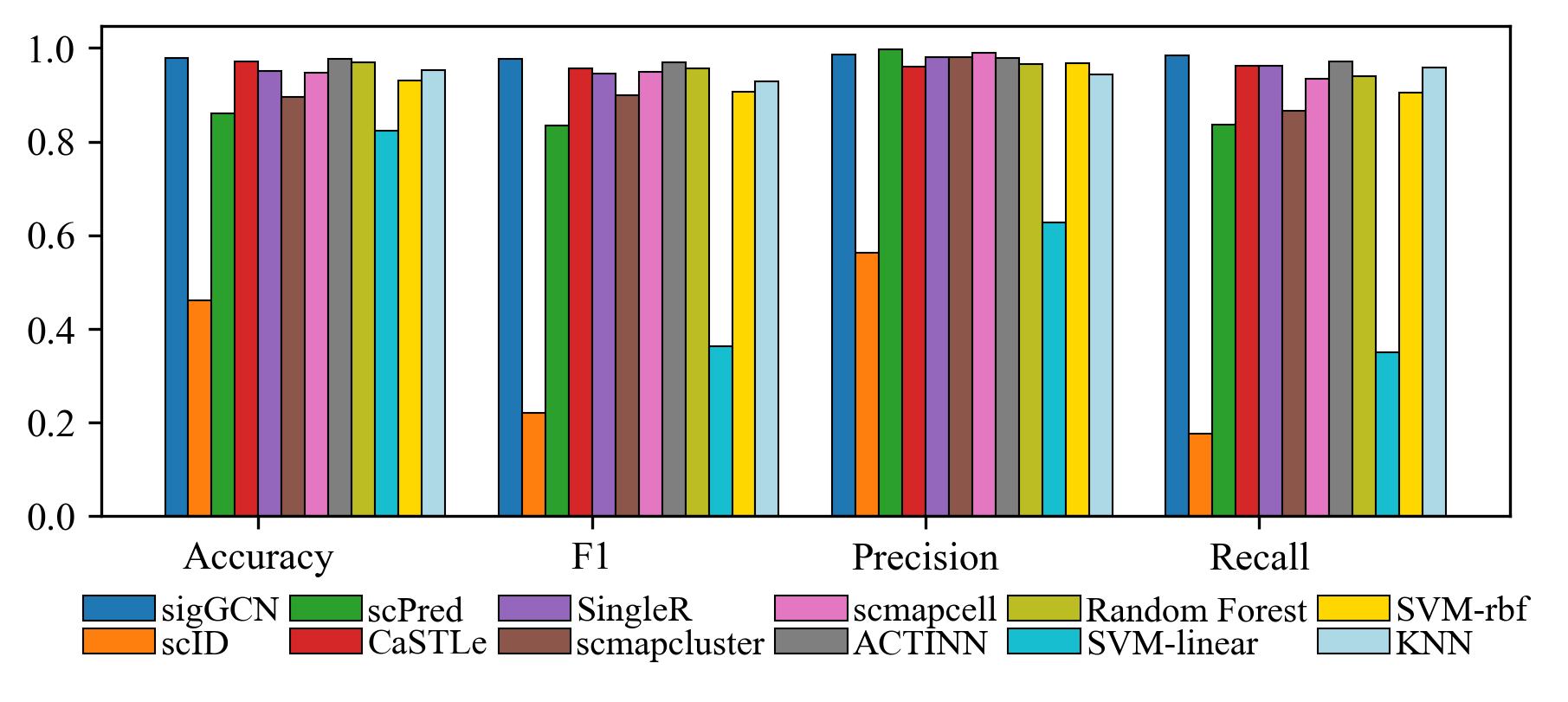

Figure S2: Bar plots of the four metrics to show the performance of scRNAseq data classifier tools and conventional classifiers on Baron Human dataset.

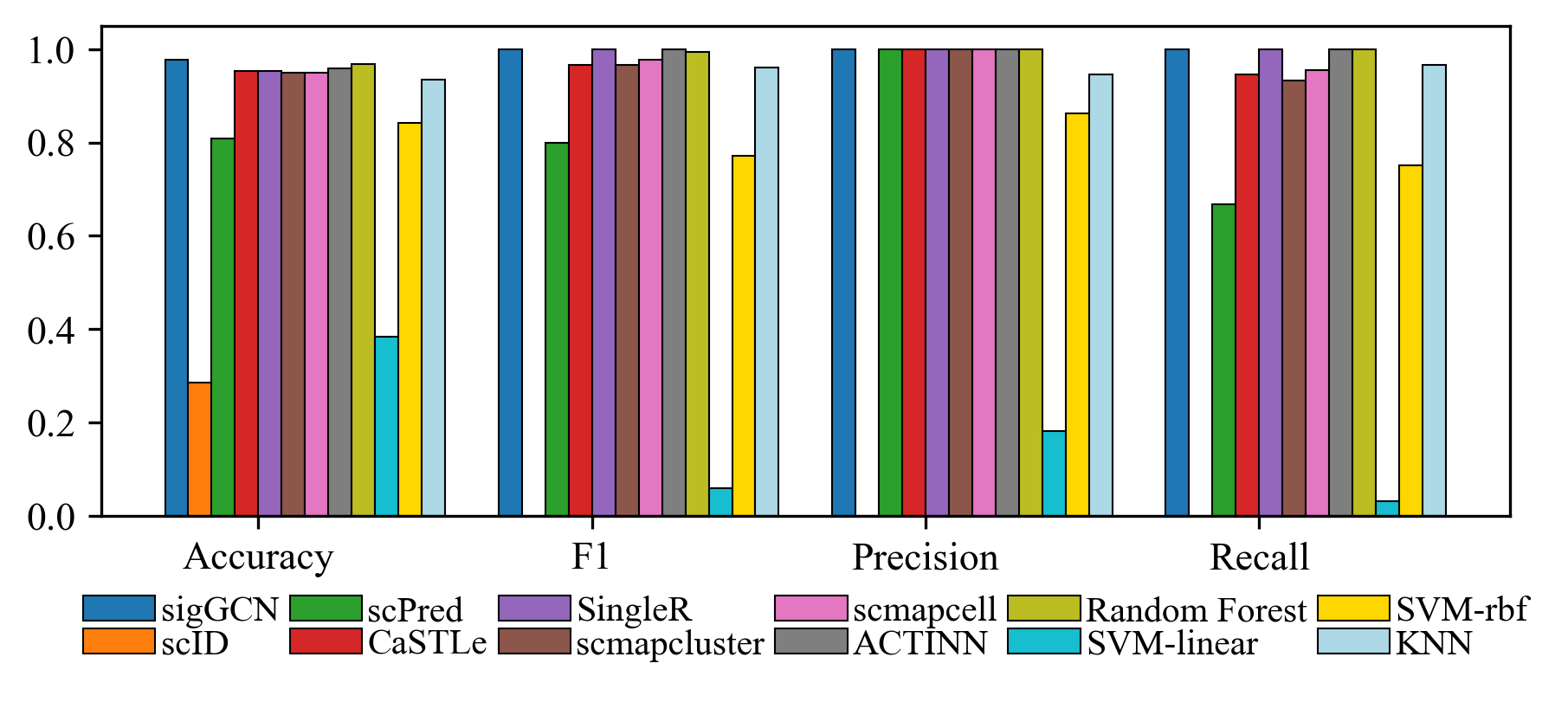

Figure S3: Bar plots of the four metrics to show the performance of scRNAseq data classifier tools and conventional classifiers on Segerstolpe dataset.

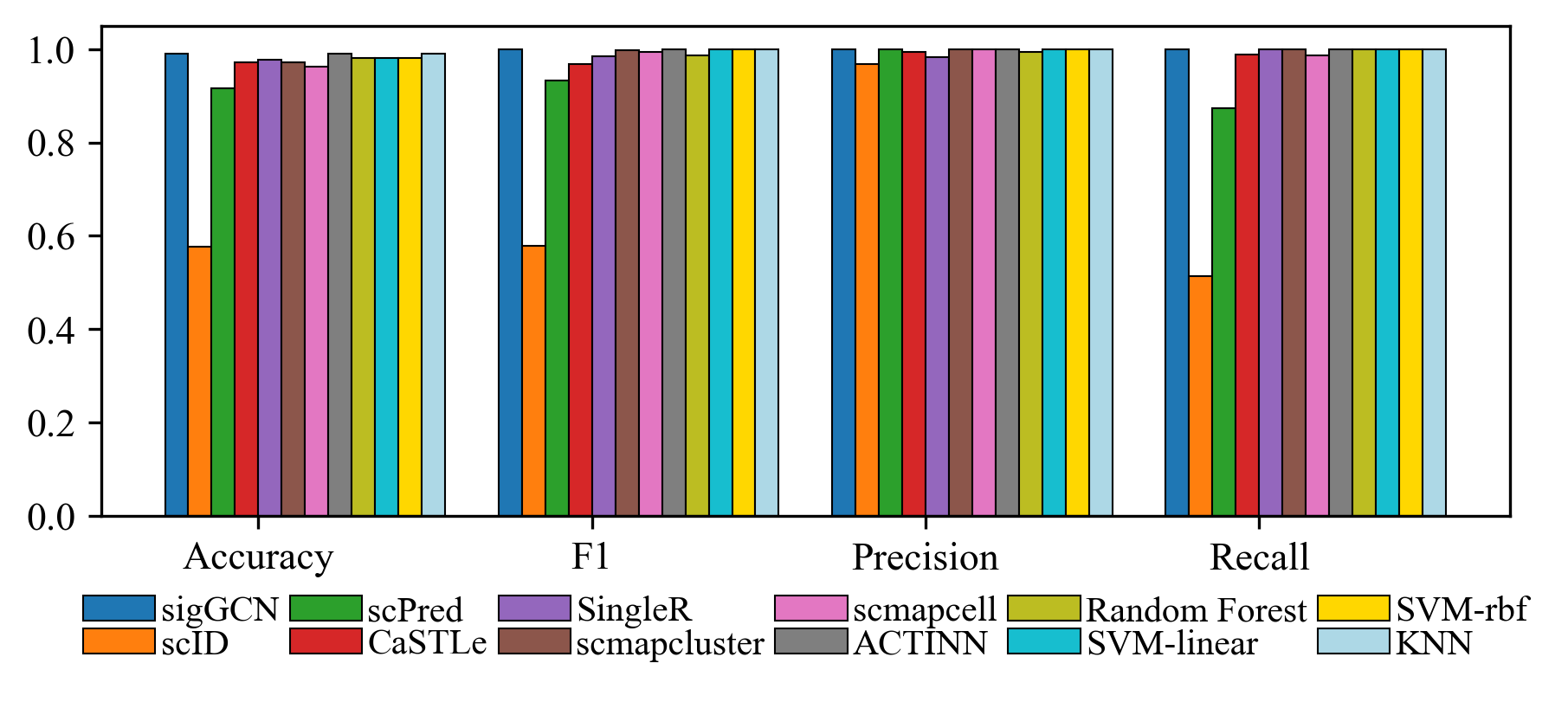

Figure S4: Bar plots of the four metrics to show the performance of scRNAseq data classifier tools and conventional classifiers on Muraro dataset.

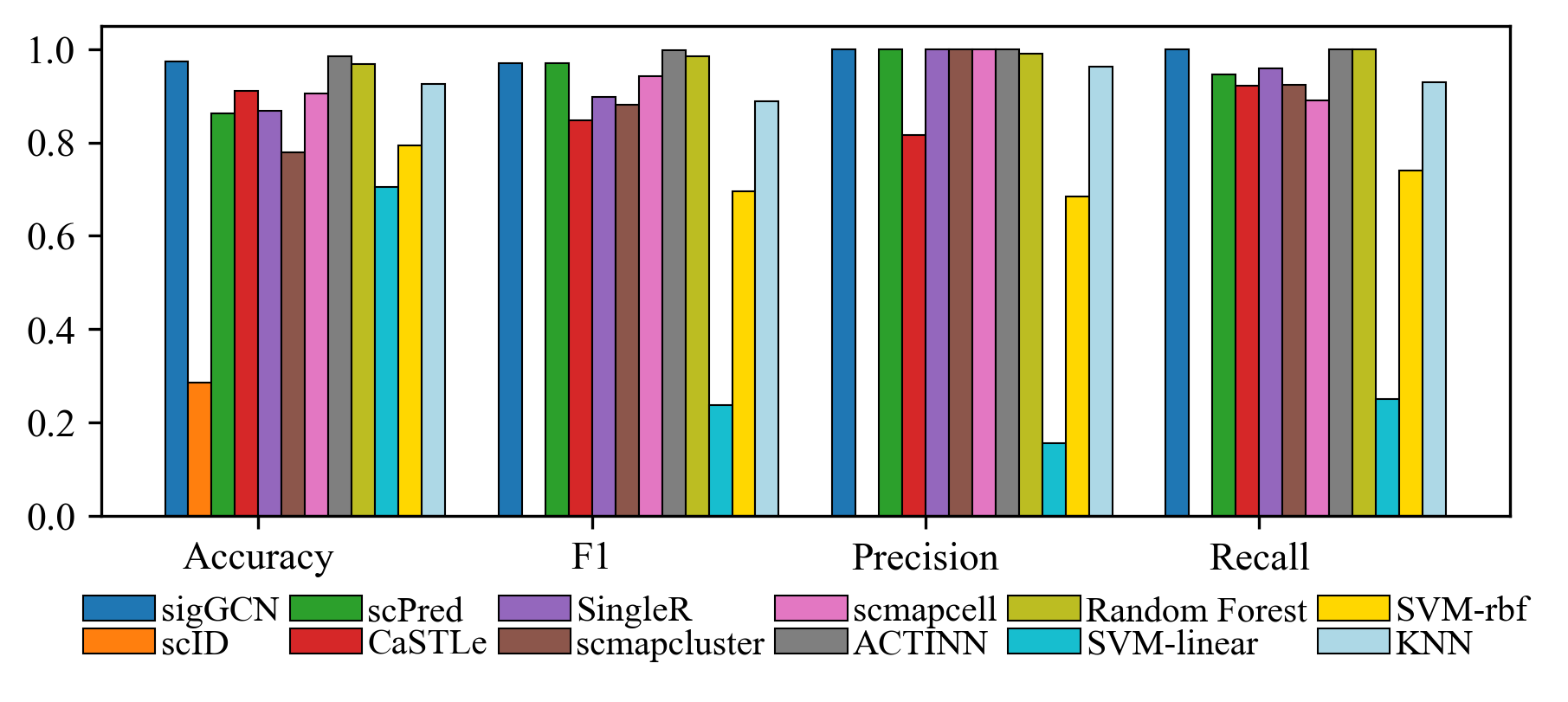

Figure S5: Bar plots of the four metrics to show the performance of scRNAseq data classifier tools and conventional classifiers on Baron Mouse dataset.

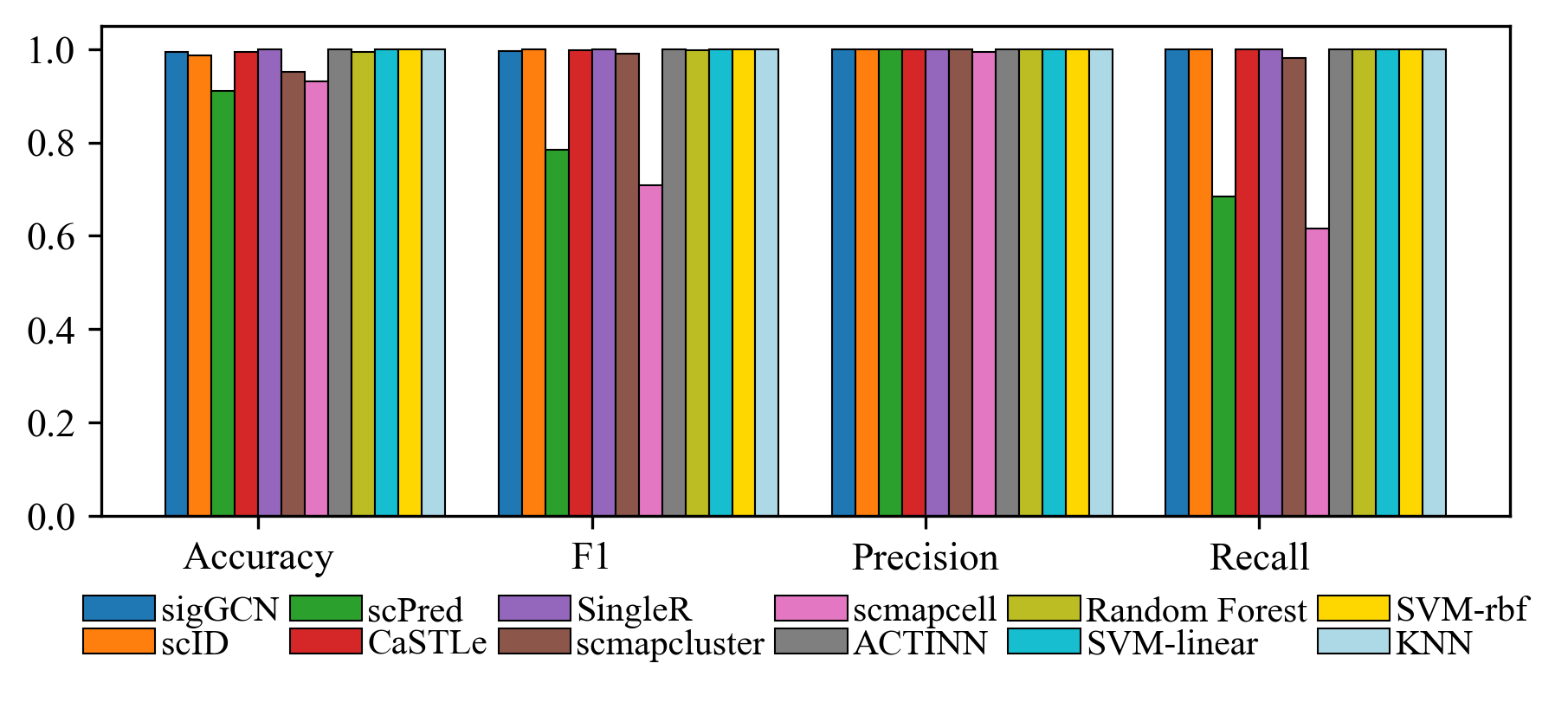

Figure S6: Bar plots of the four metrics to show the performance of scRNAseq data classifier tools and conventional classifiers on Xin dataset.

(a)	(b)

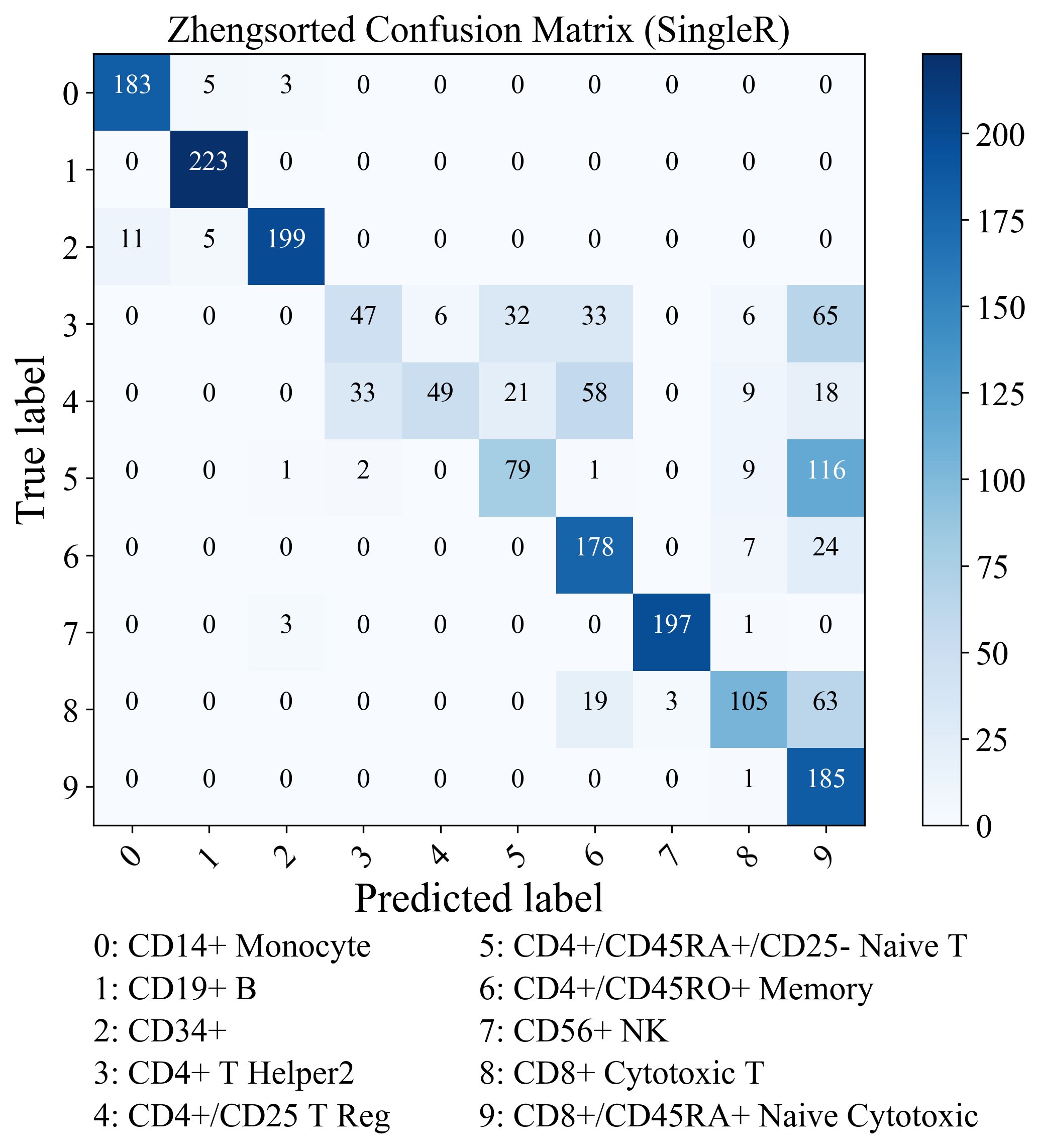

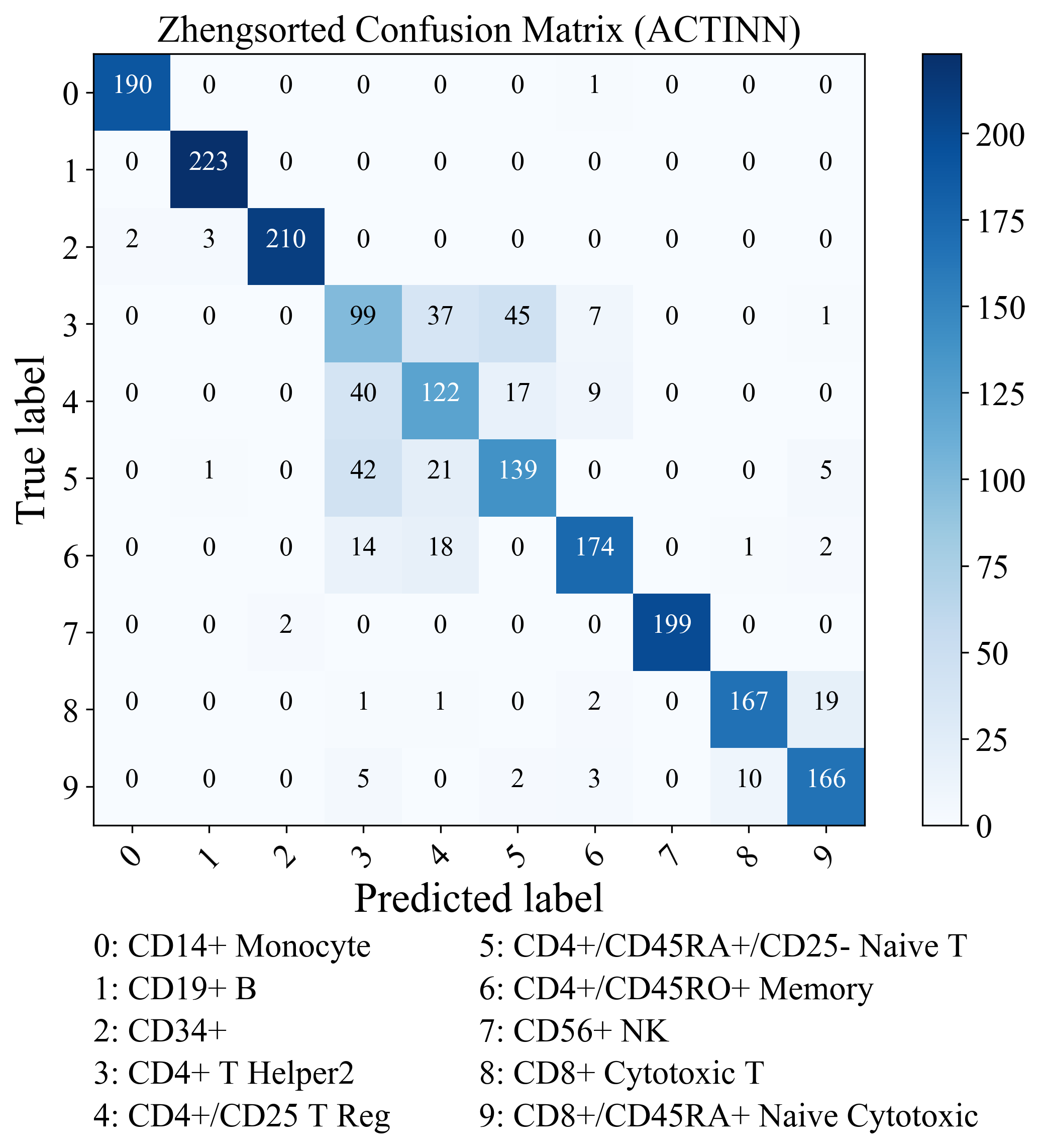

(c)

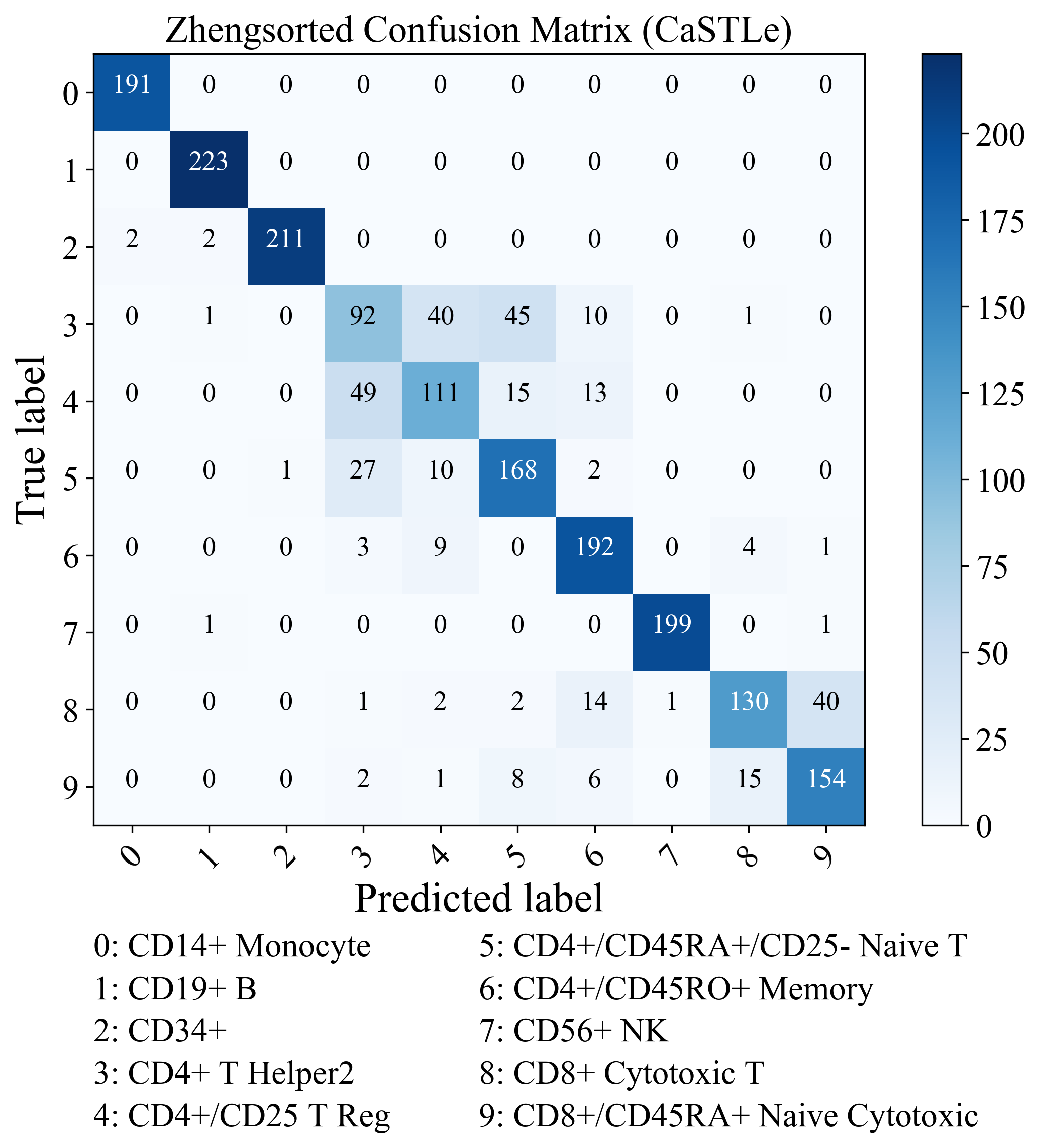

Figure S7: Confusion matrix of the class predictions using (a) SingleR, (b) ACTINN, (c) CasTLe on the Zhengsorted dataset.

(a)	(b)

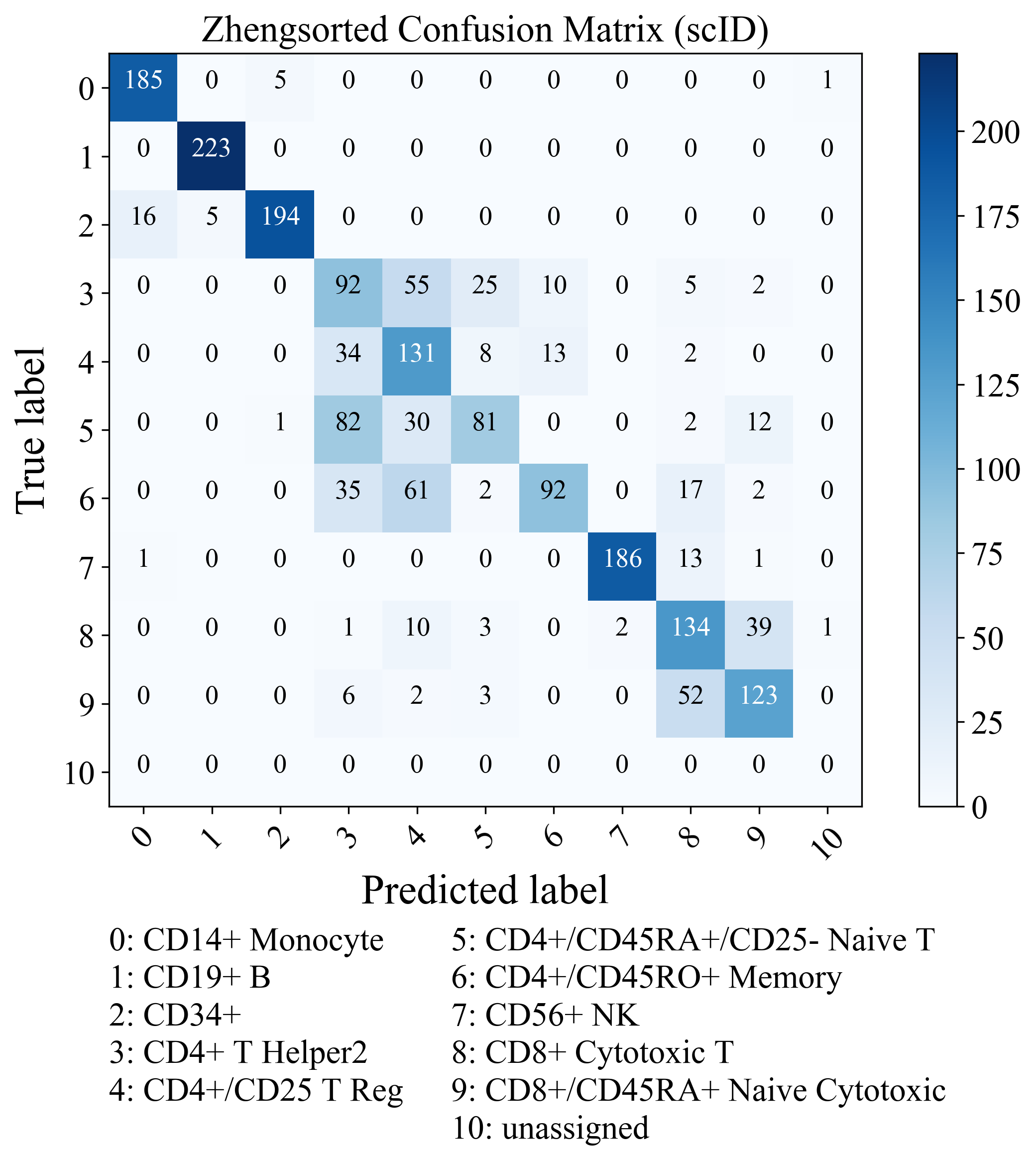

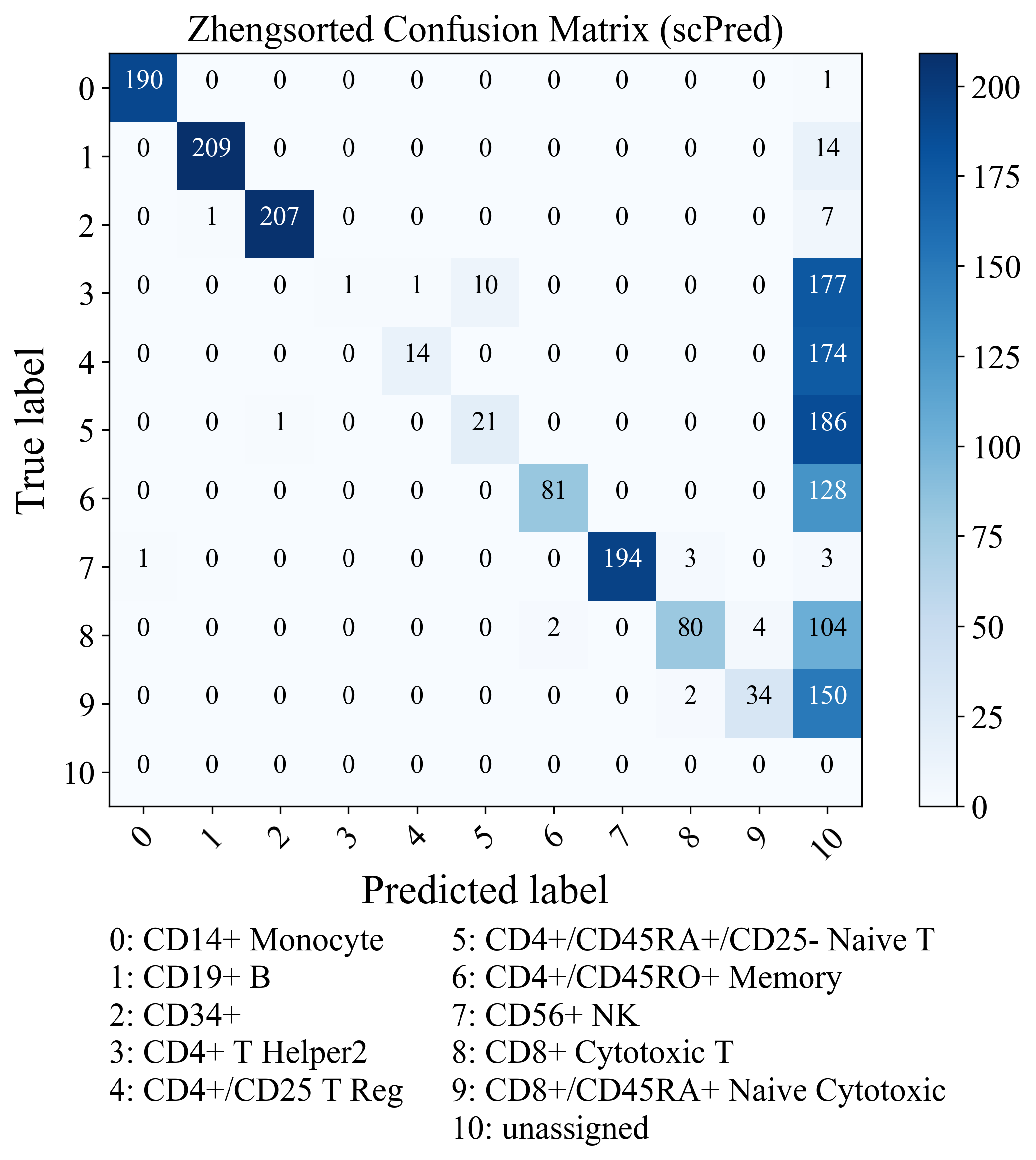

(c)	(d)

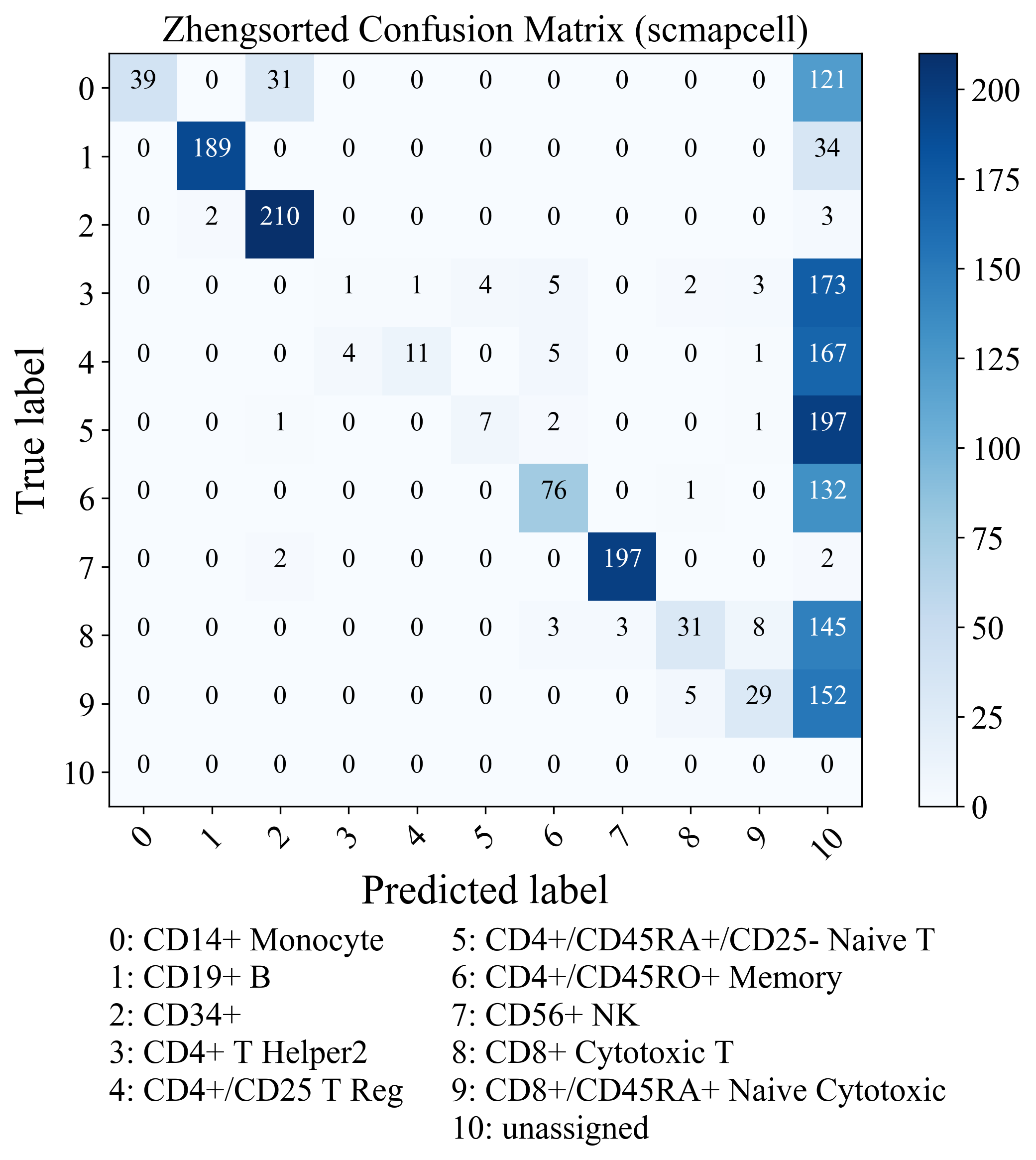

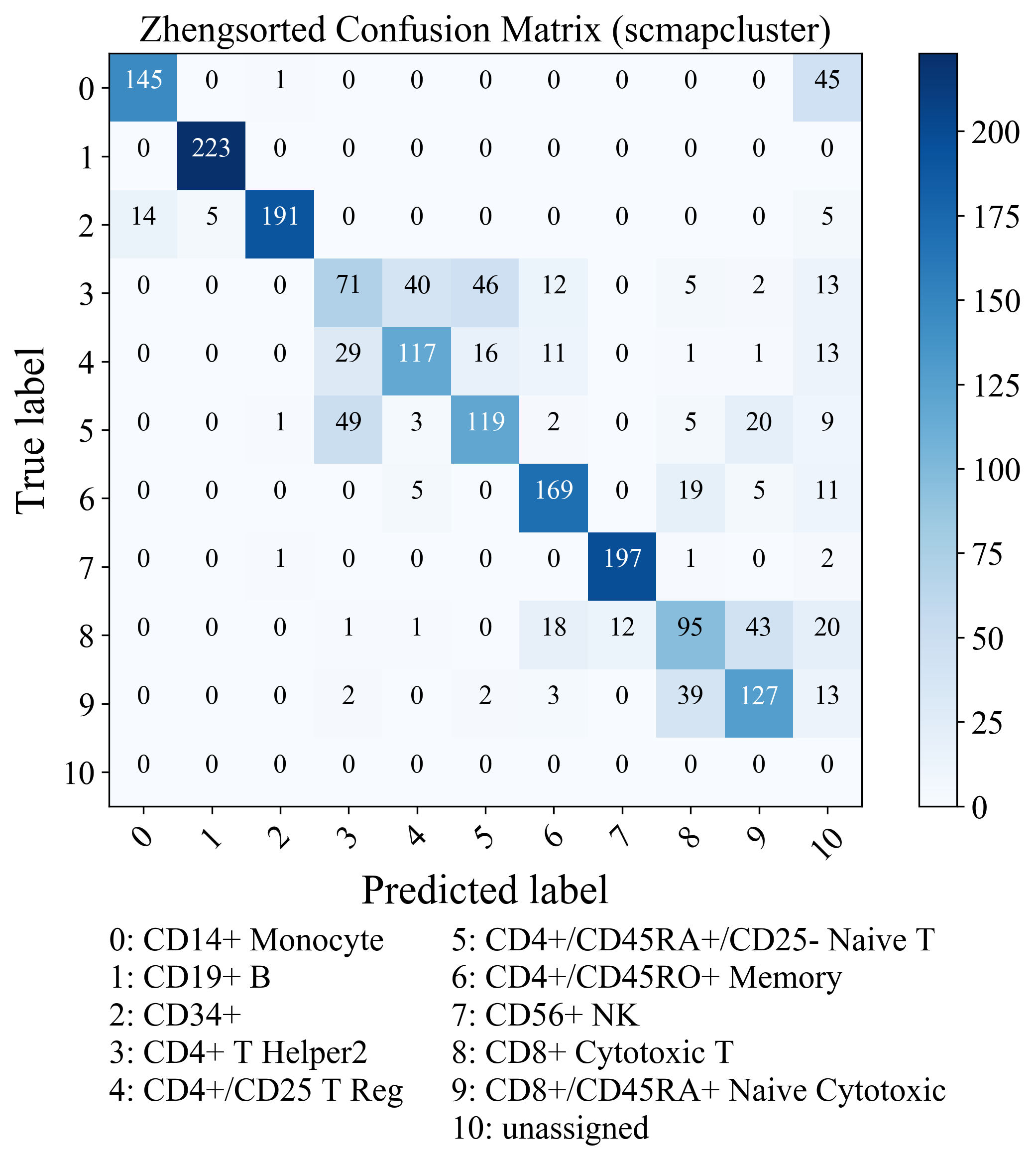

Figure S8: Confusion matrix of the class predictions using (a) scID, (b) scPred, (c) scmapcell, (d) scmapcluster on the Zhengsorted dataset.

(a)	(b)

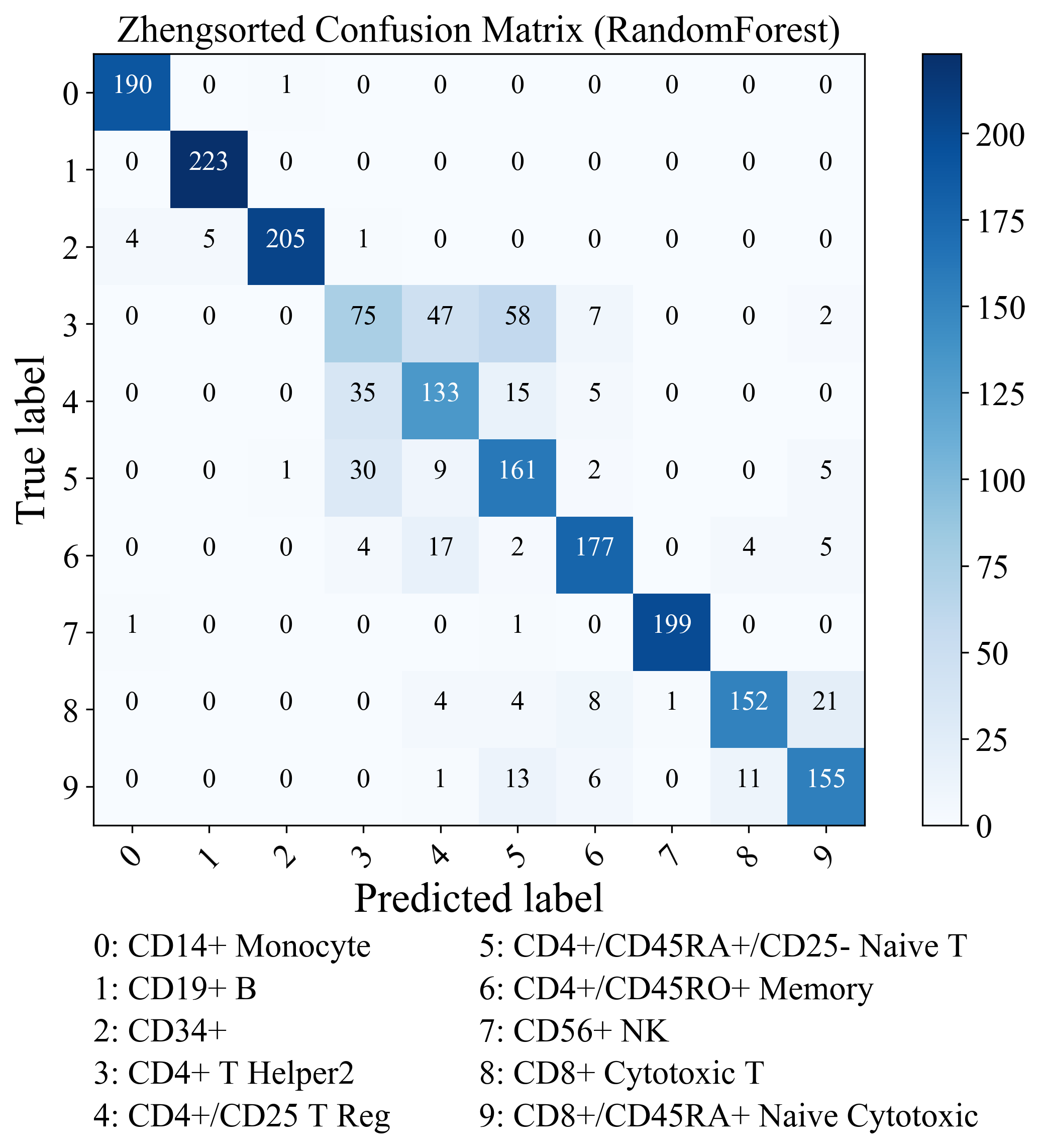

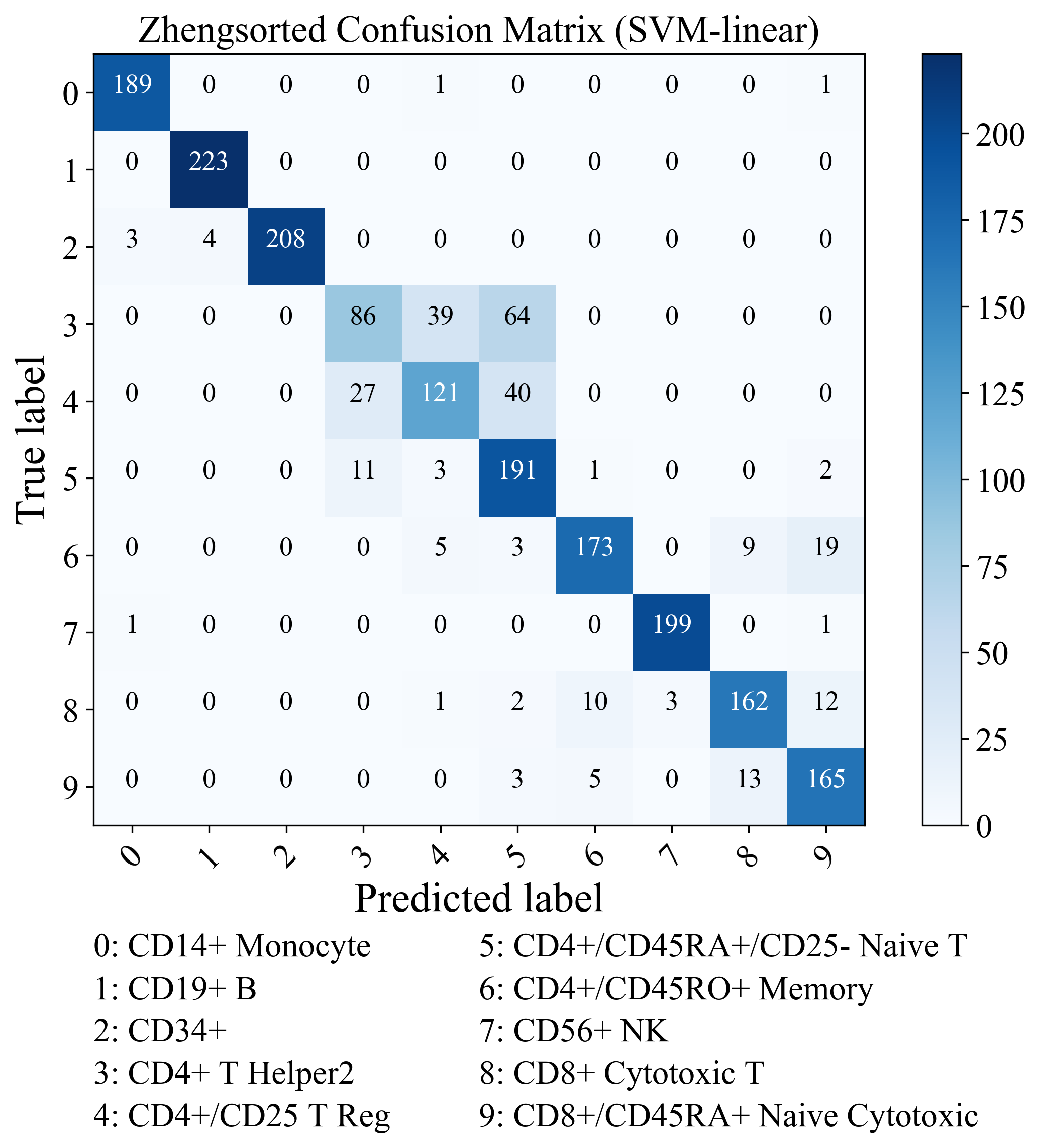

(c)	(d)

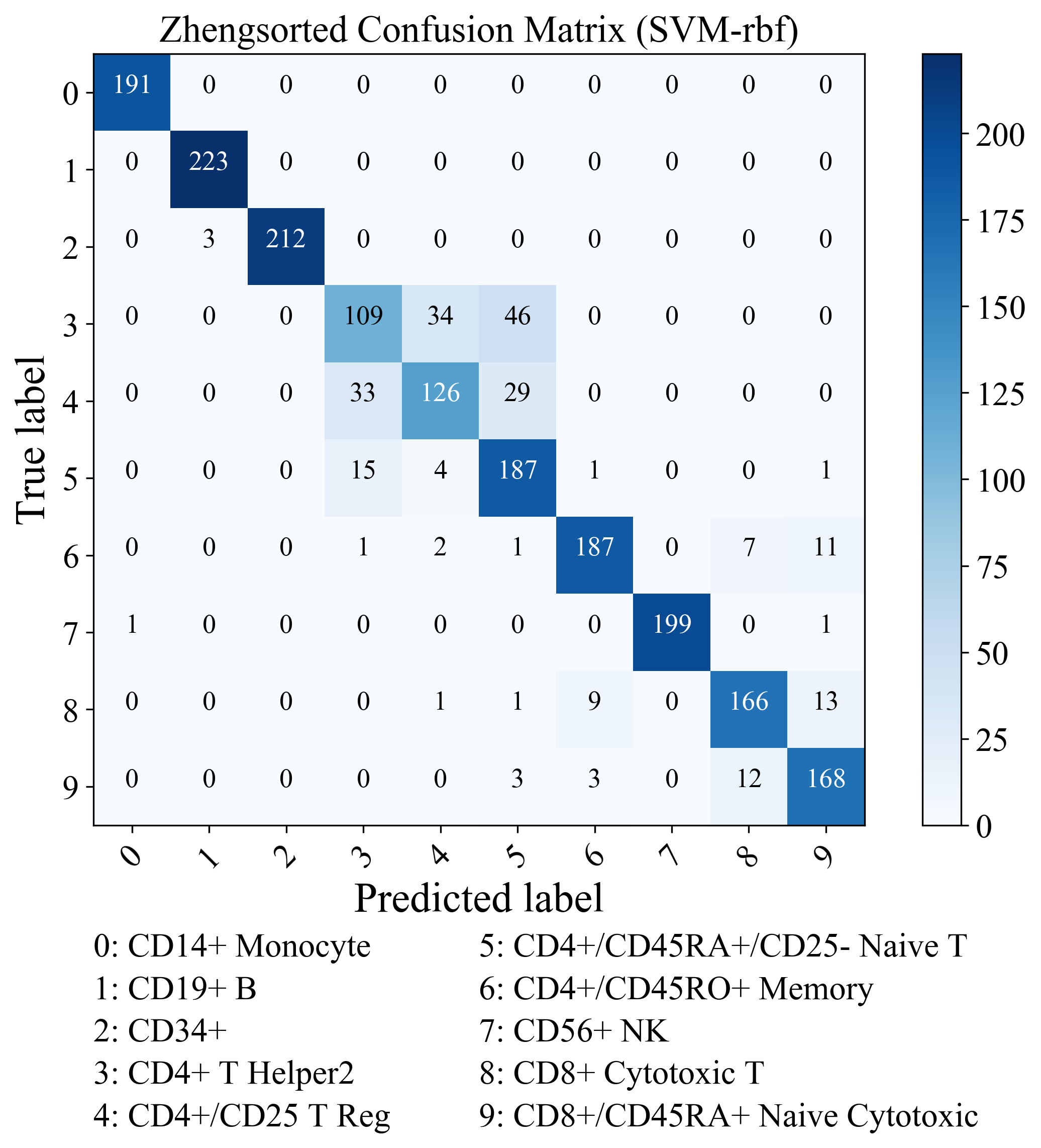

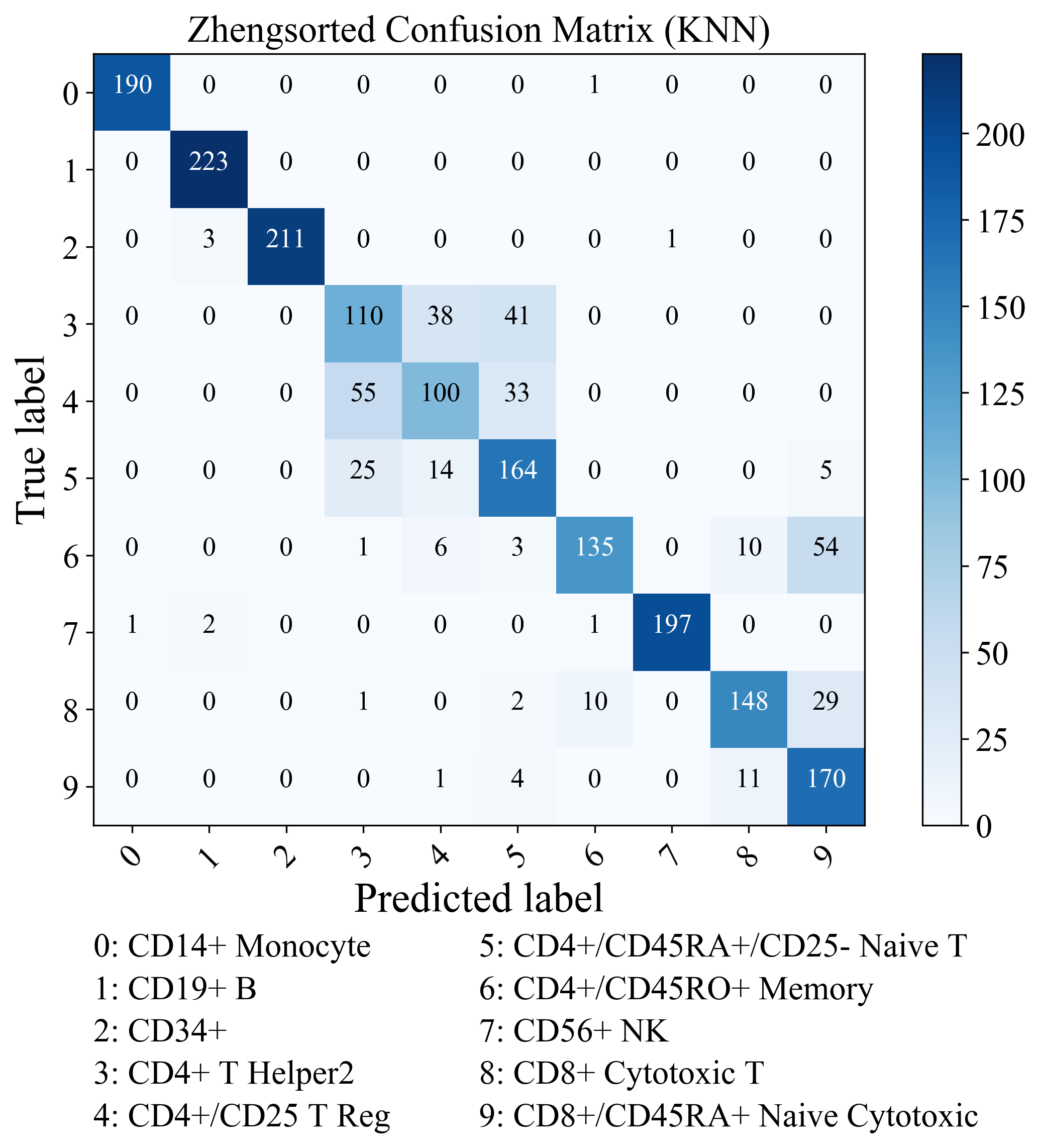

Figure S9: Confusion matrix of the class predictions using (a) Random Forest, (b) SVM-linear, (c) SVM-rbf, (d) KNN on the Zhengsorted dataset.

(a)	(b)

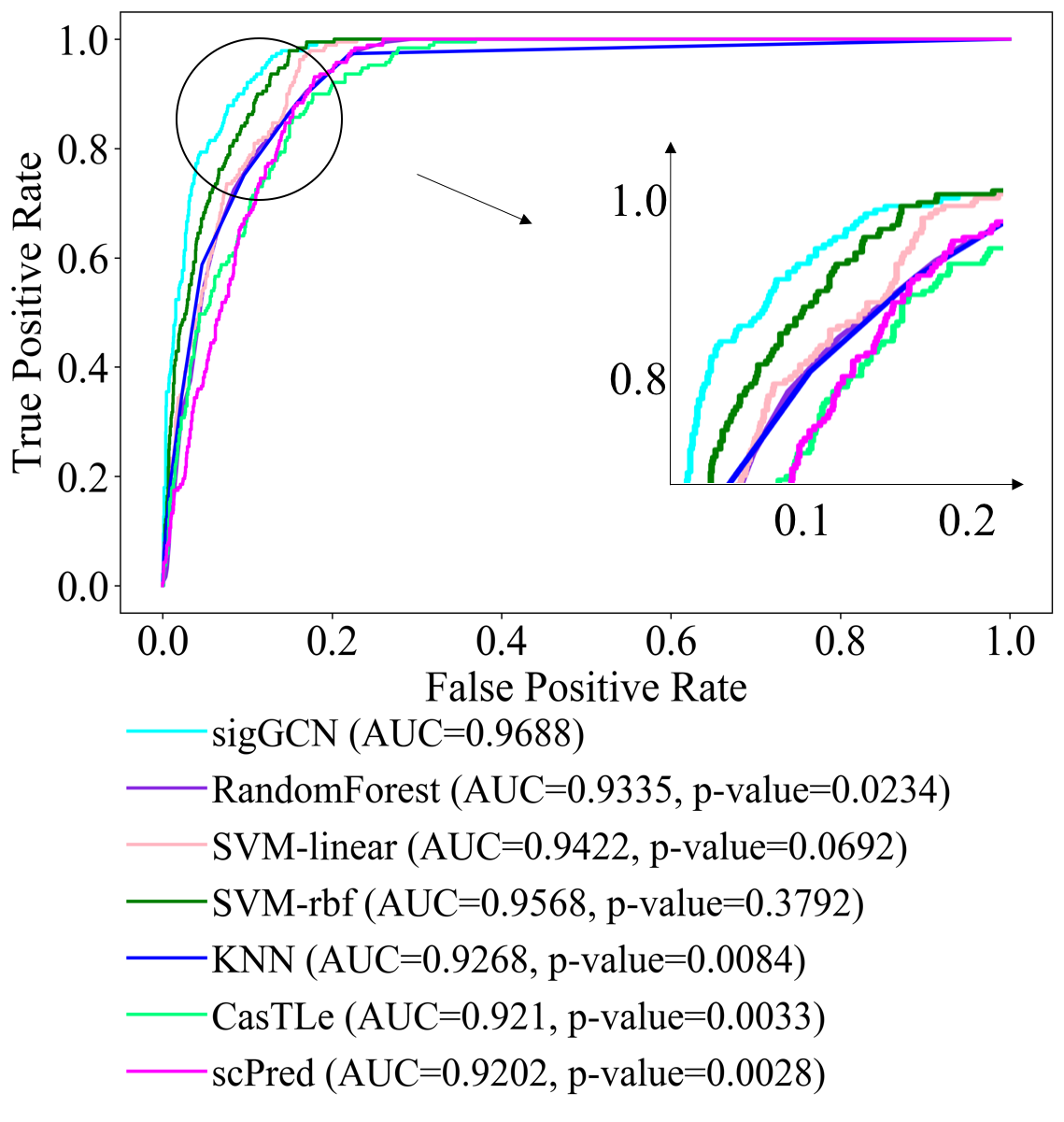

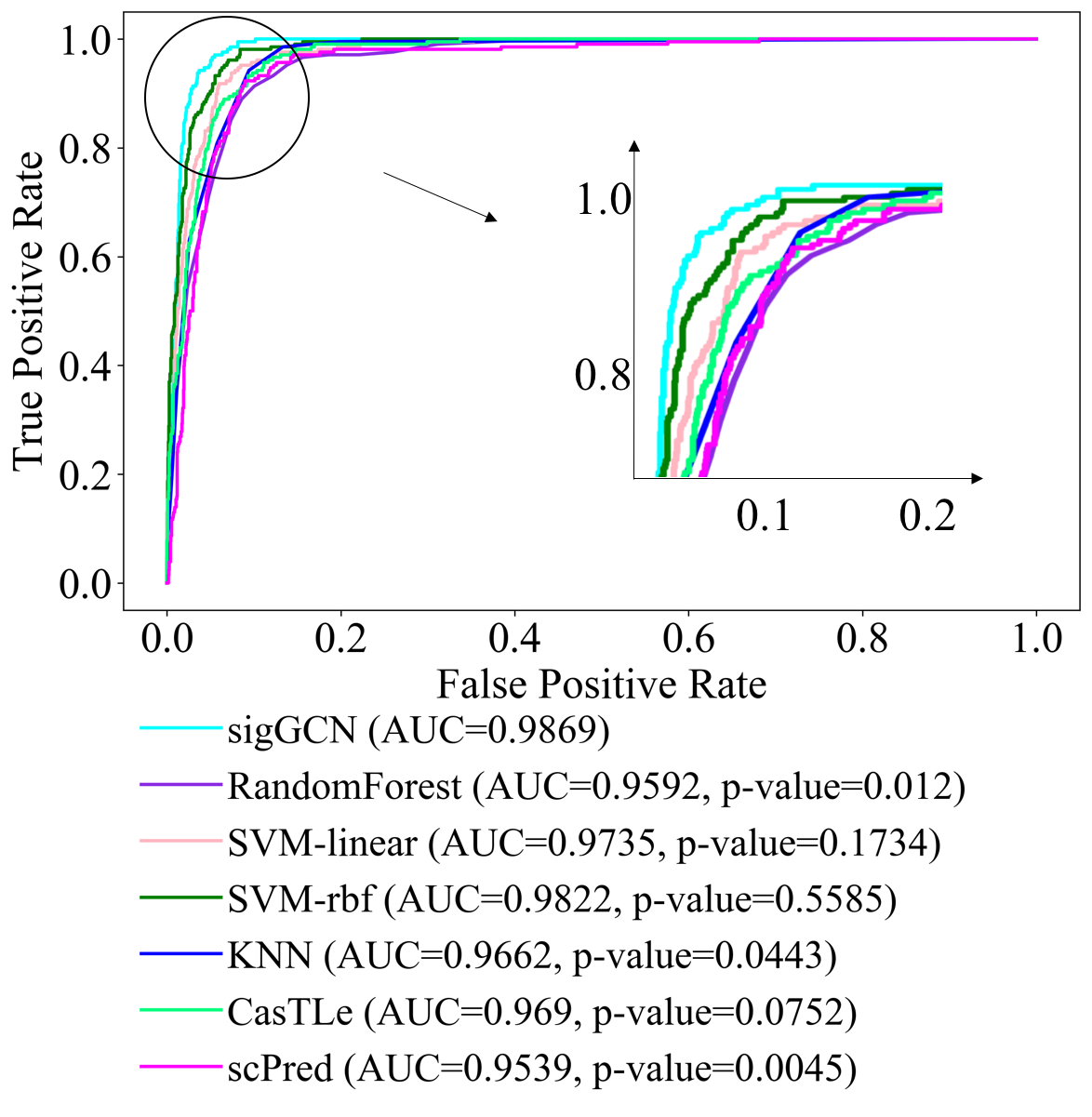

(c)	(d)

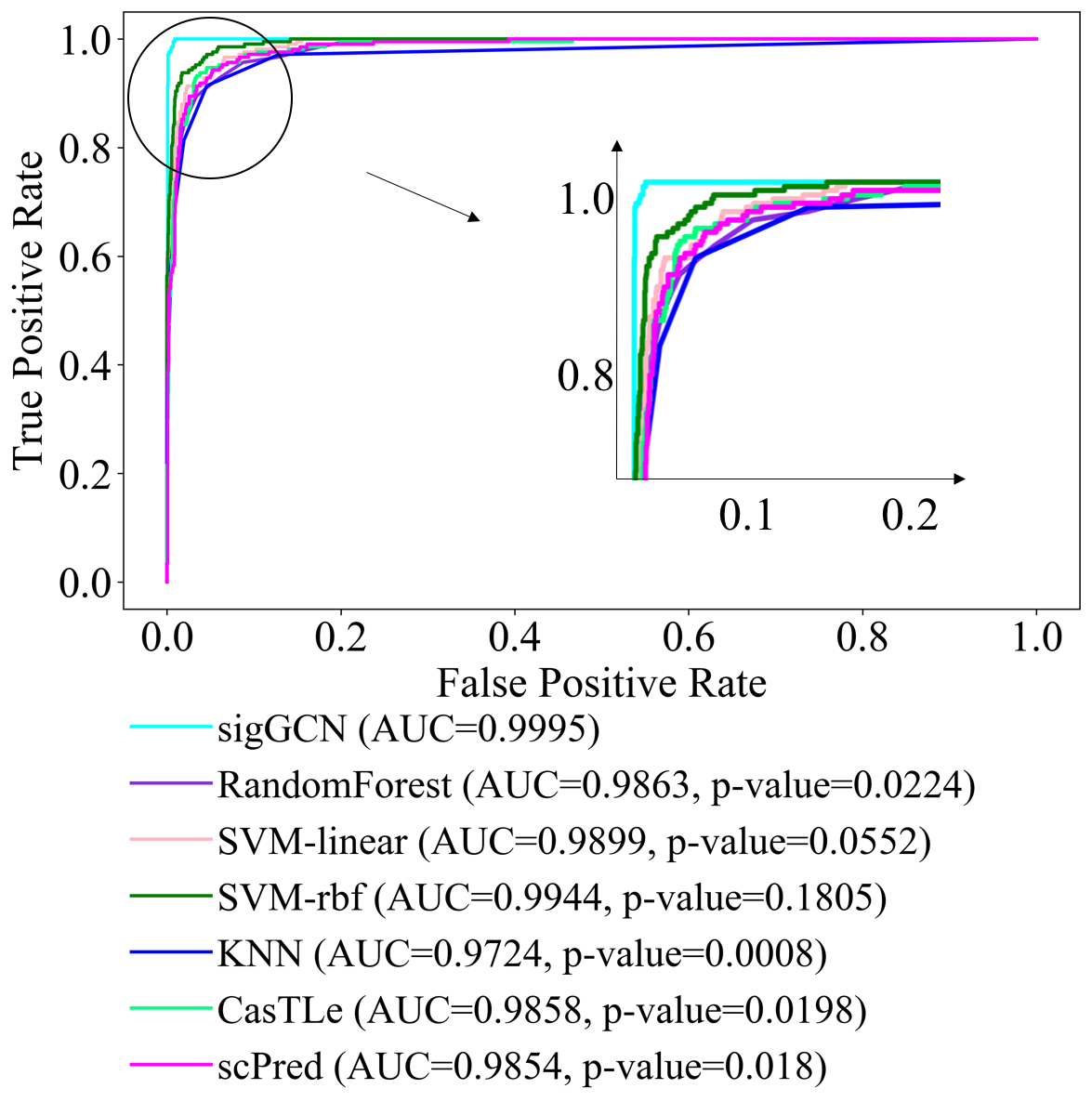

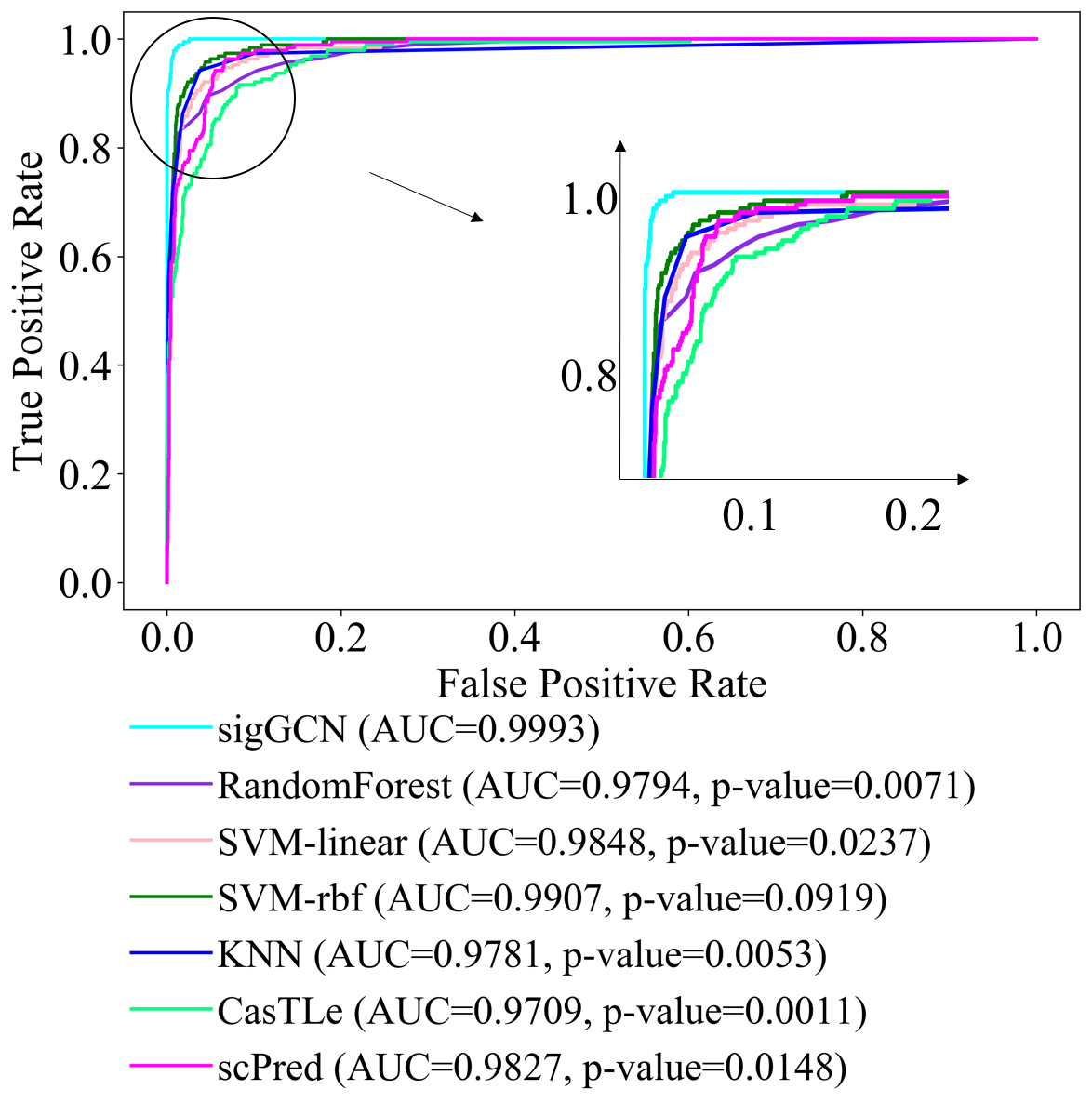

Figure S10: ROC curves and the p-values calculated by McNeil & Hanley's test that show the significance of difference between the areas under the ROC curve of sigGCN and that of each method. (a) ROC curves of class 3 (CD4+ T Helper) and the p-values, (b) ROC curves of class 5 (CD4+/CD45RA+/CD25- Naive T) and the p-values, (c) ROC curves of class 6 (CD4+/CD45RO+ Memory) and the p-values, and (d) ROC curves of class 8 (CD8+ Cytotoxic T) and the p-values.

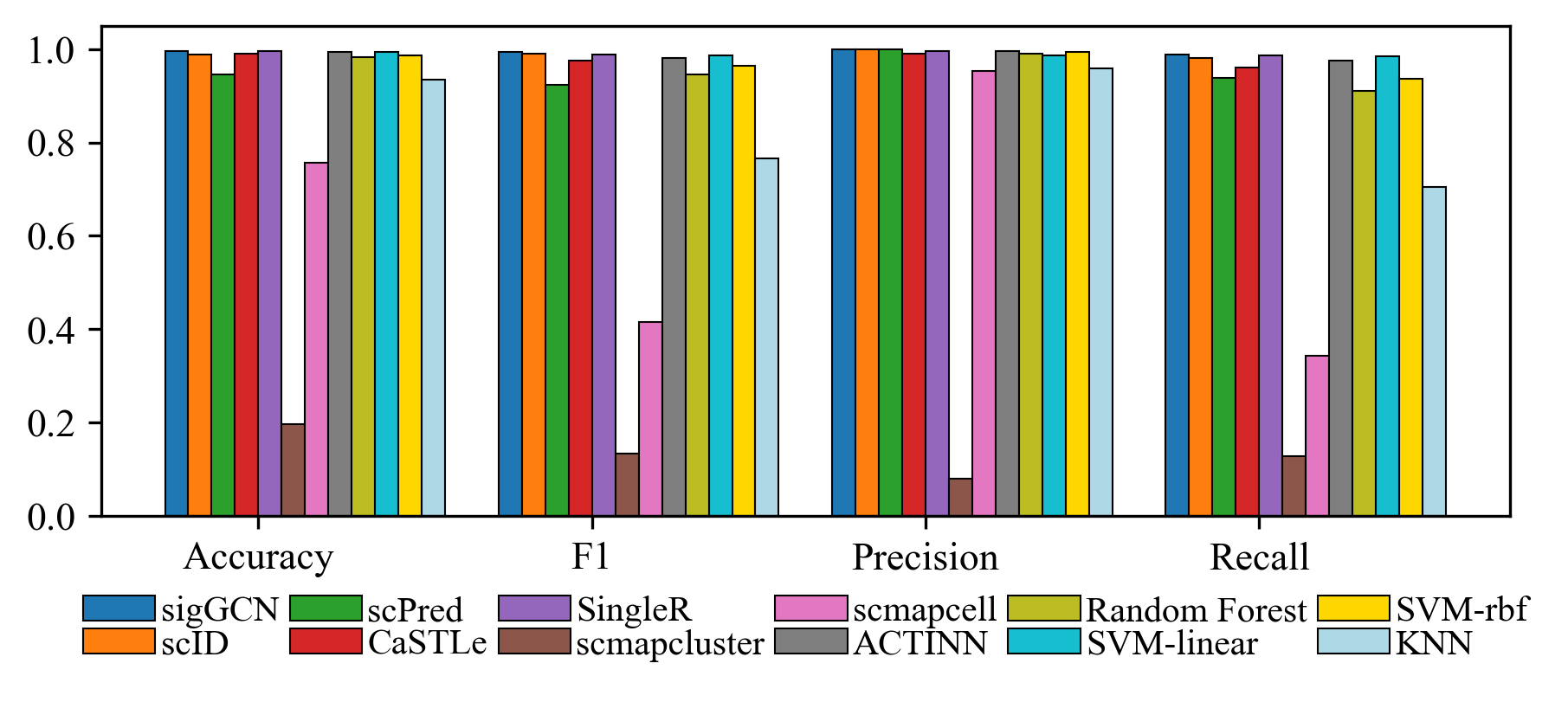

Figure S11: Bar plots of the four metrics to show the performance of scRNAseq data classifier tools and conventional classifiers when training on Baron Human, Muraro, Segerstolpe and testing on Xin dataset.

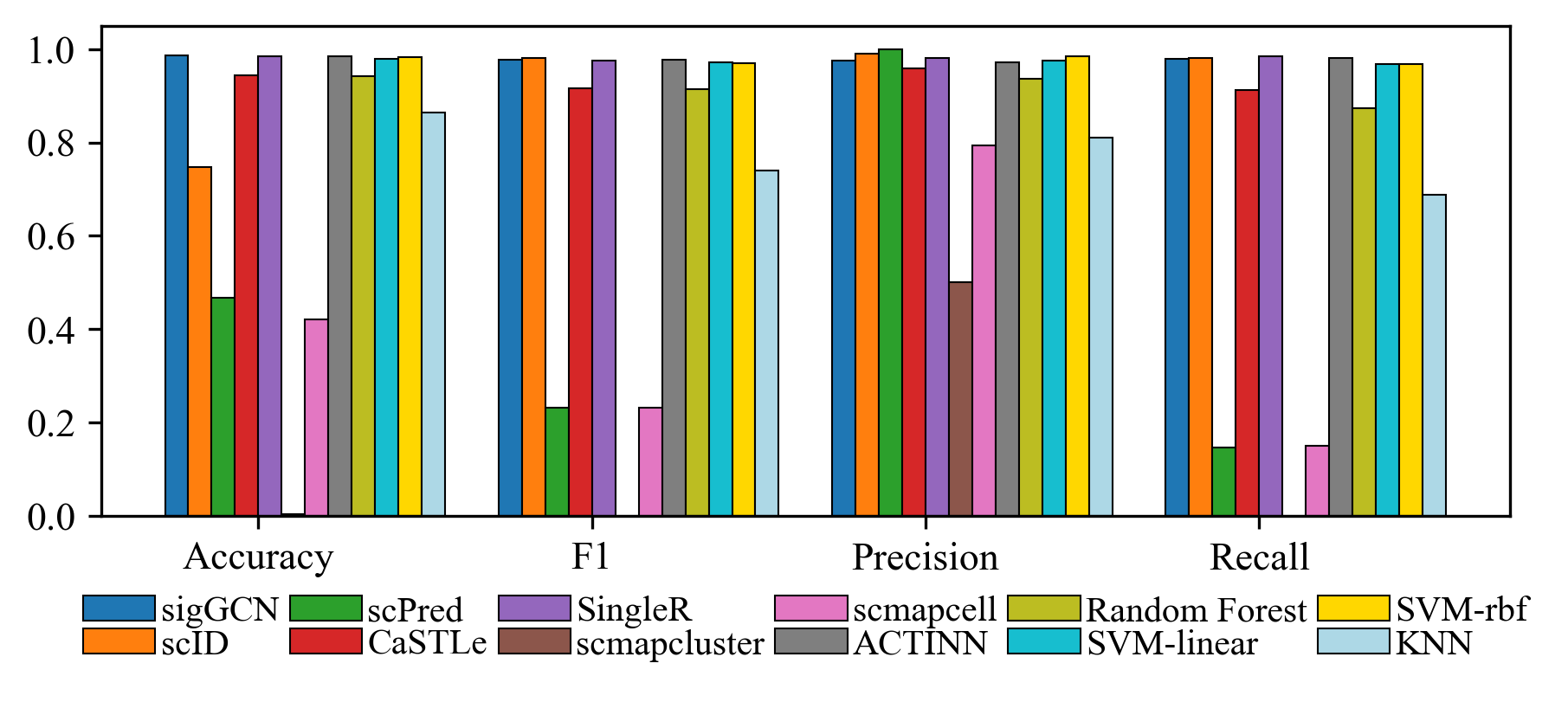

Figure S12: Bar plots of the four metrics to show the performance of scRNAseq data classifier tools and conventional classifiers when training on Xin, Muraro, Segerstolpe and testing on Baron Human dataset.

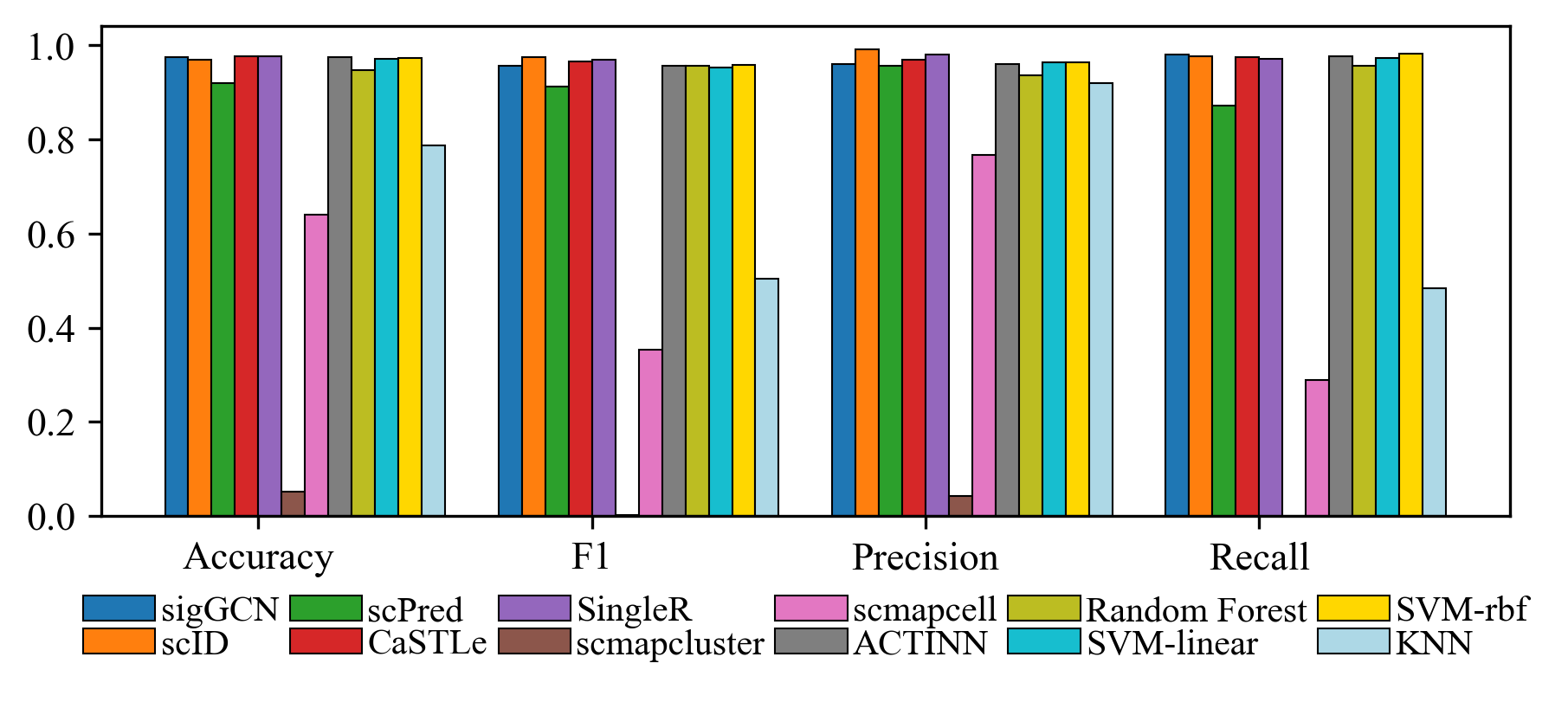

Figure S13: Bar plots of the four metrics to show the performance of scRNAseq data classifier tools and conventional classifiers when training on Xin, Baron Human, Segerstolpe and testing on Muraro dataset.

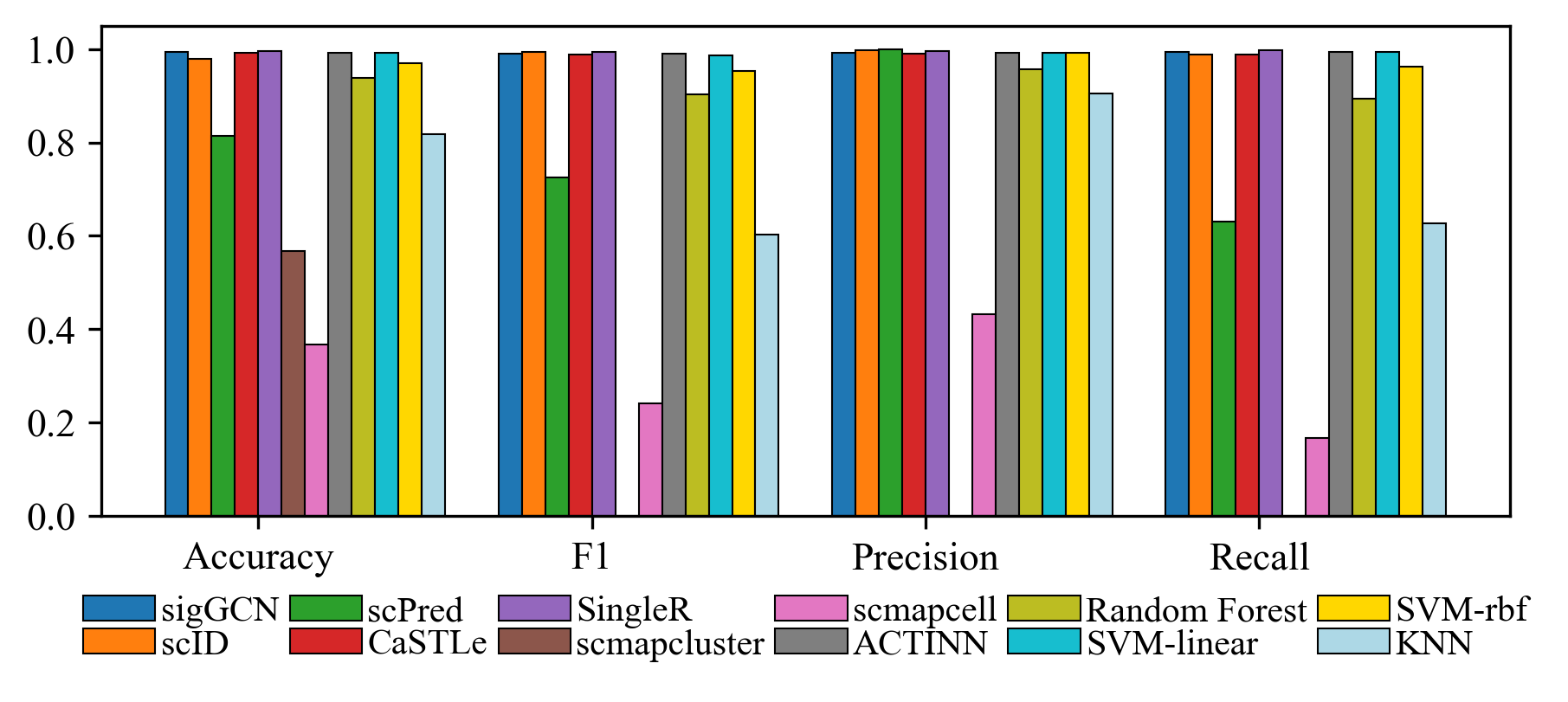

Figure S14: Bar plots of the four metrics to show the performance of scRNAseq data classifier tools and conventional classifiers when training on Xin, Baron Human, Muraro and testing on Segerstolpe dataset.

(a)	(b)	(c)

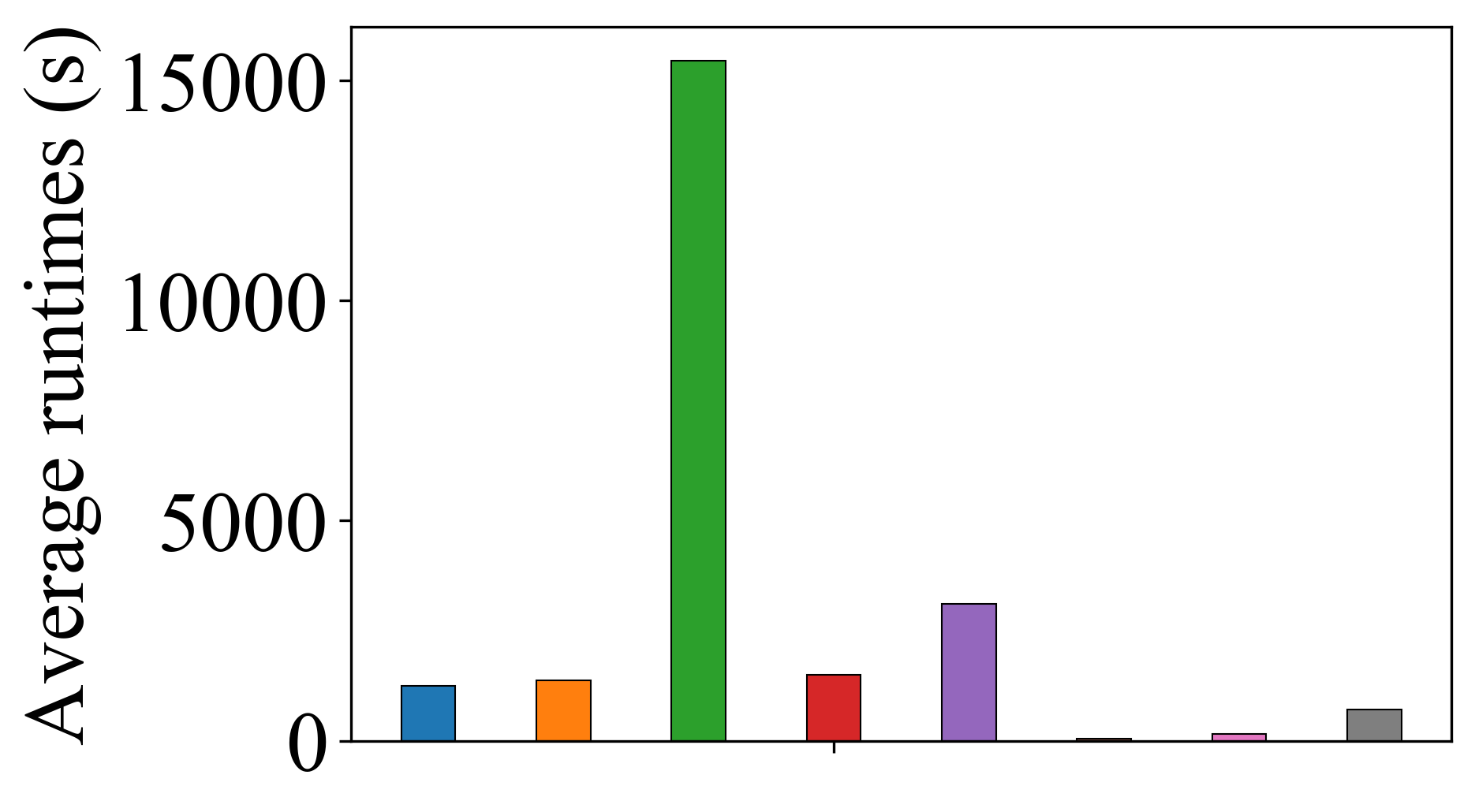

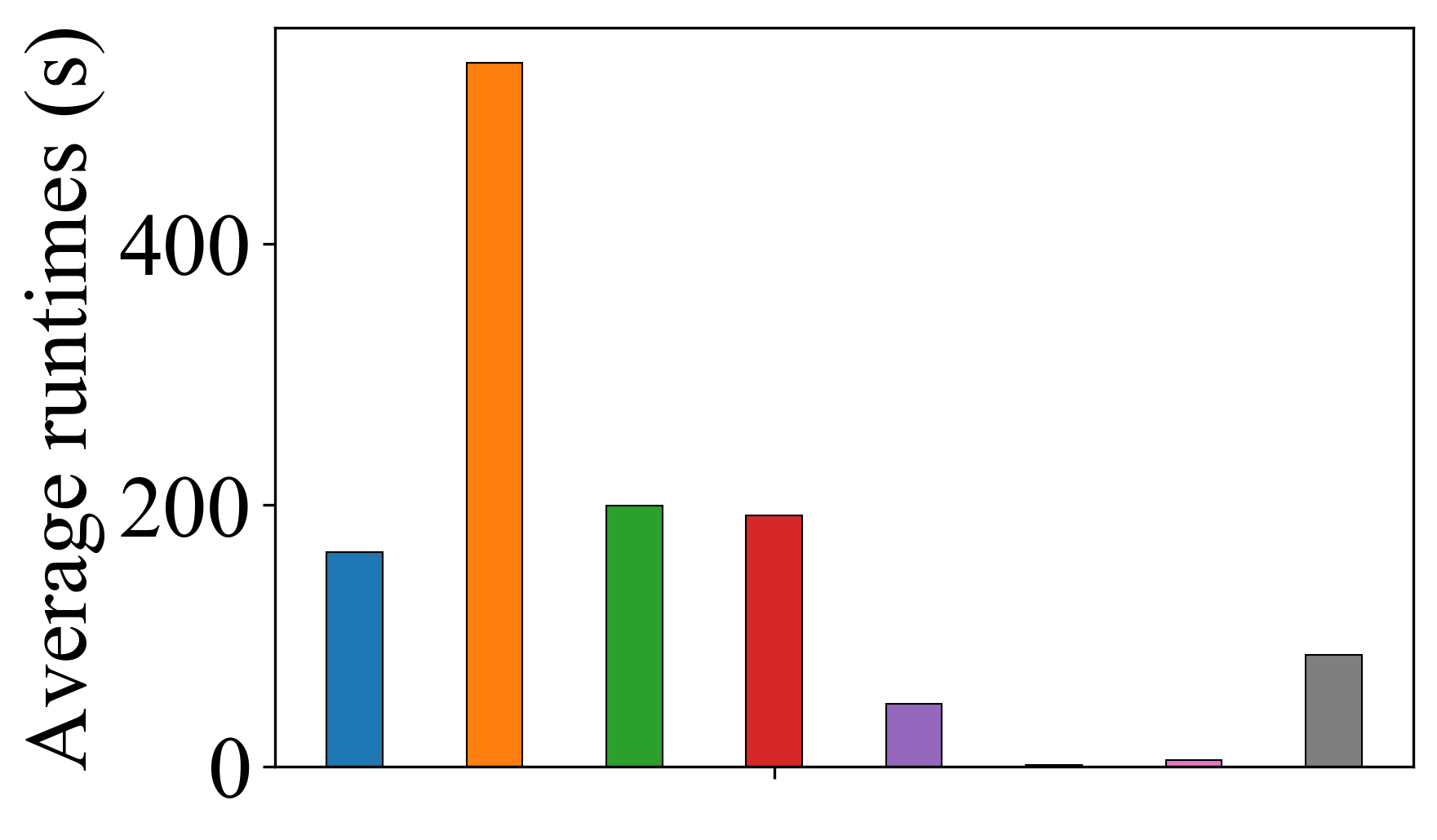

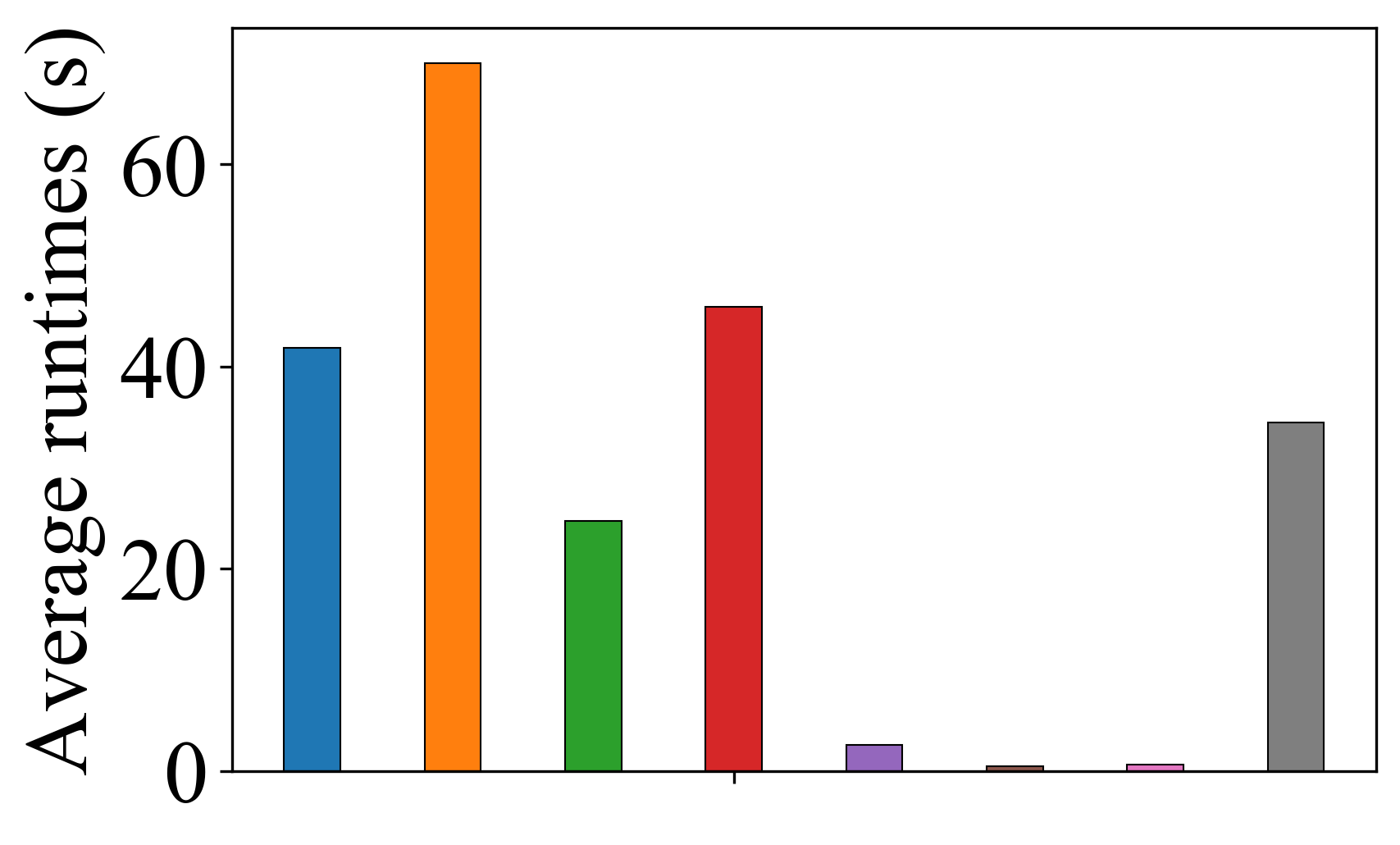

(d)	(e)	(f)

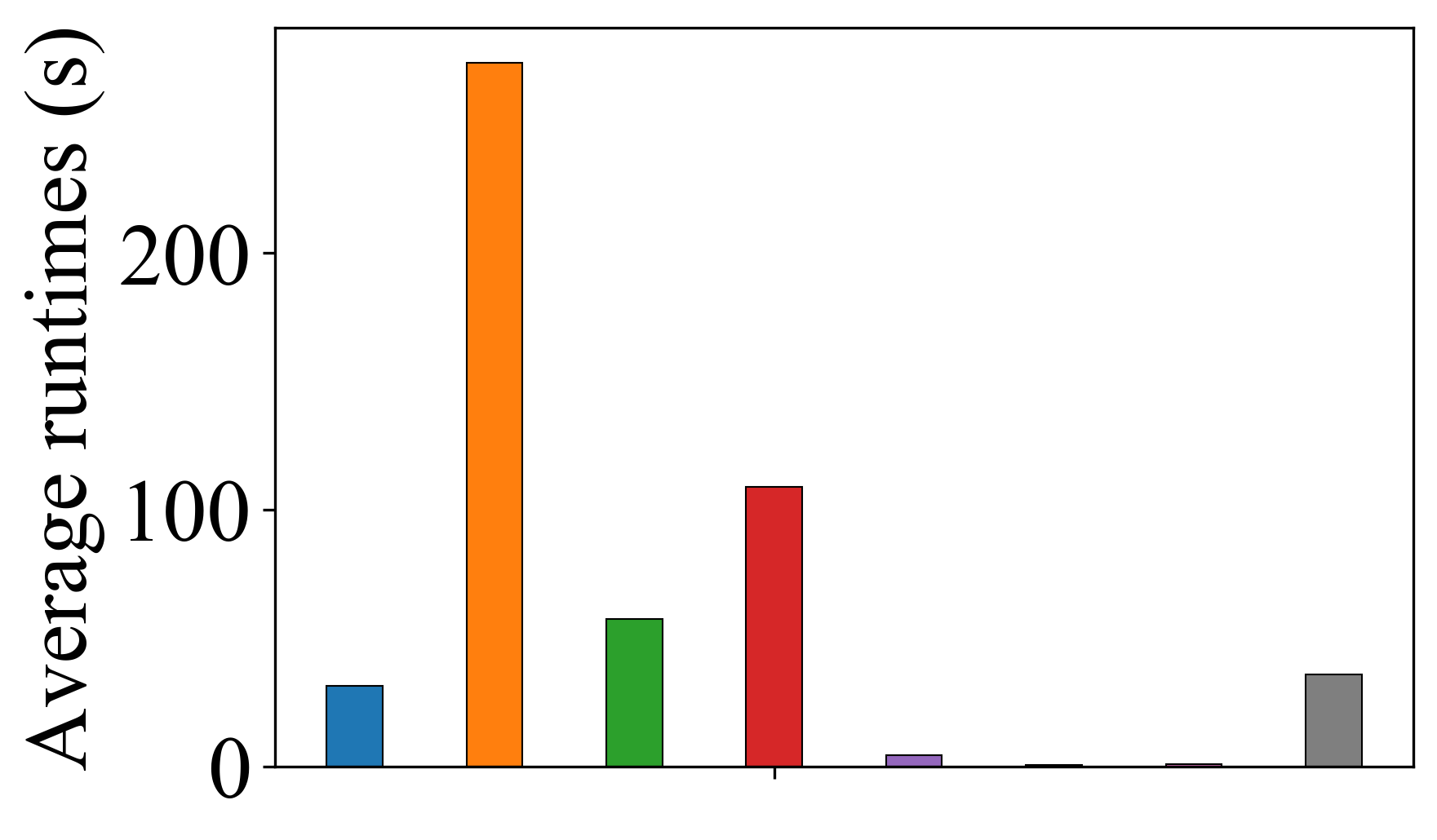

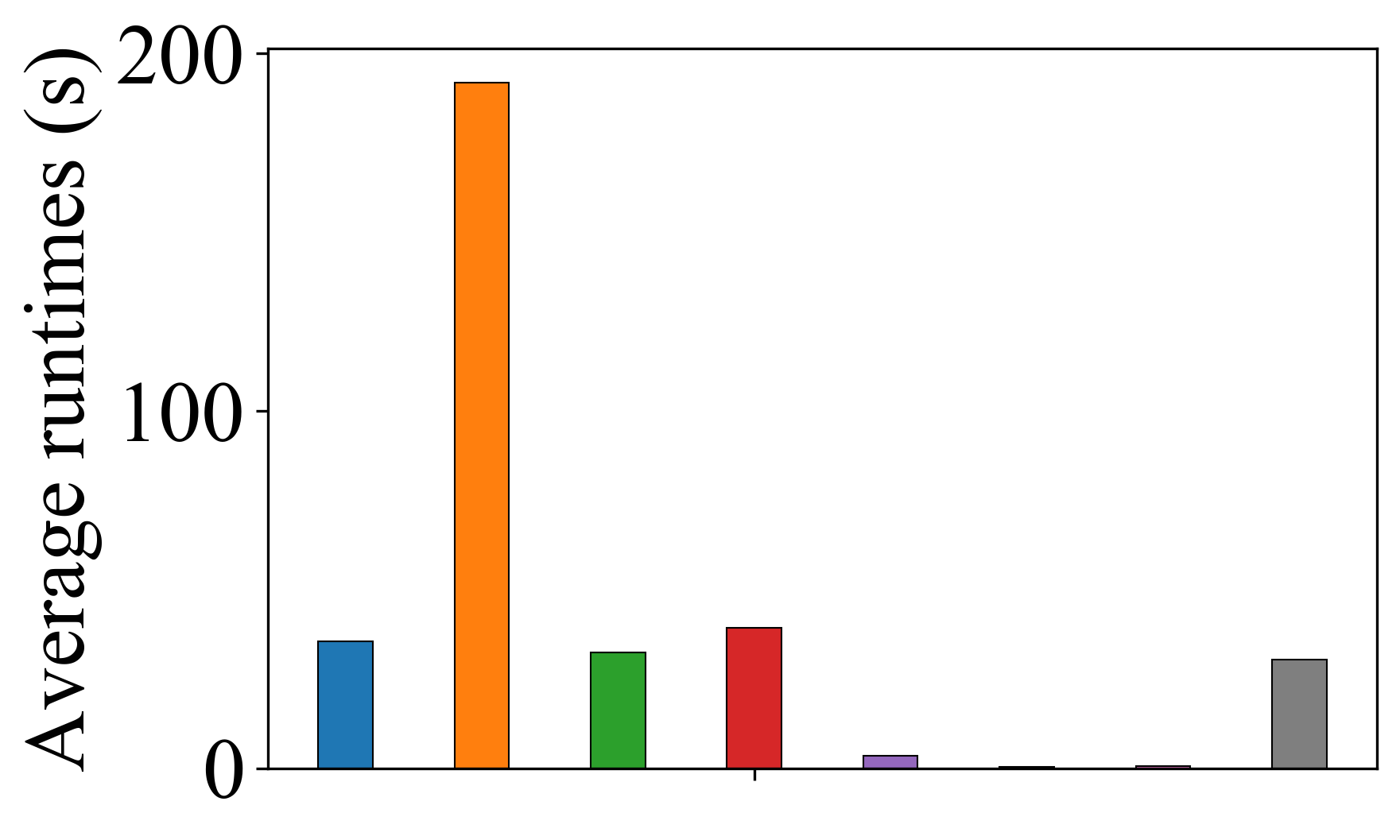

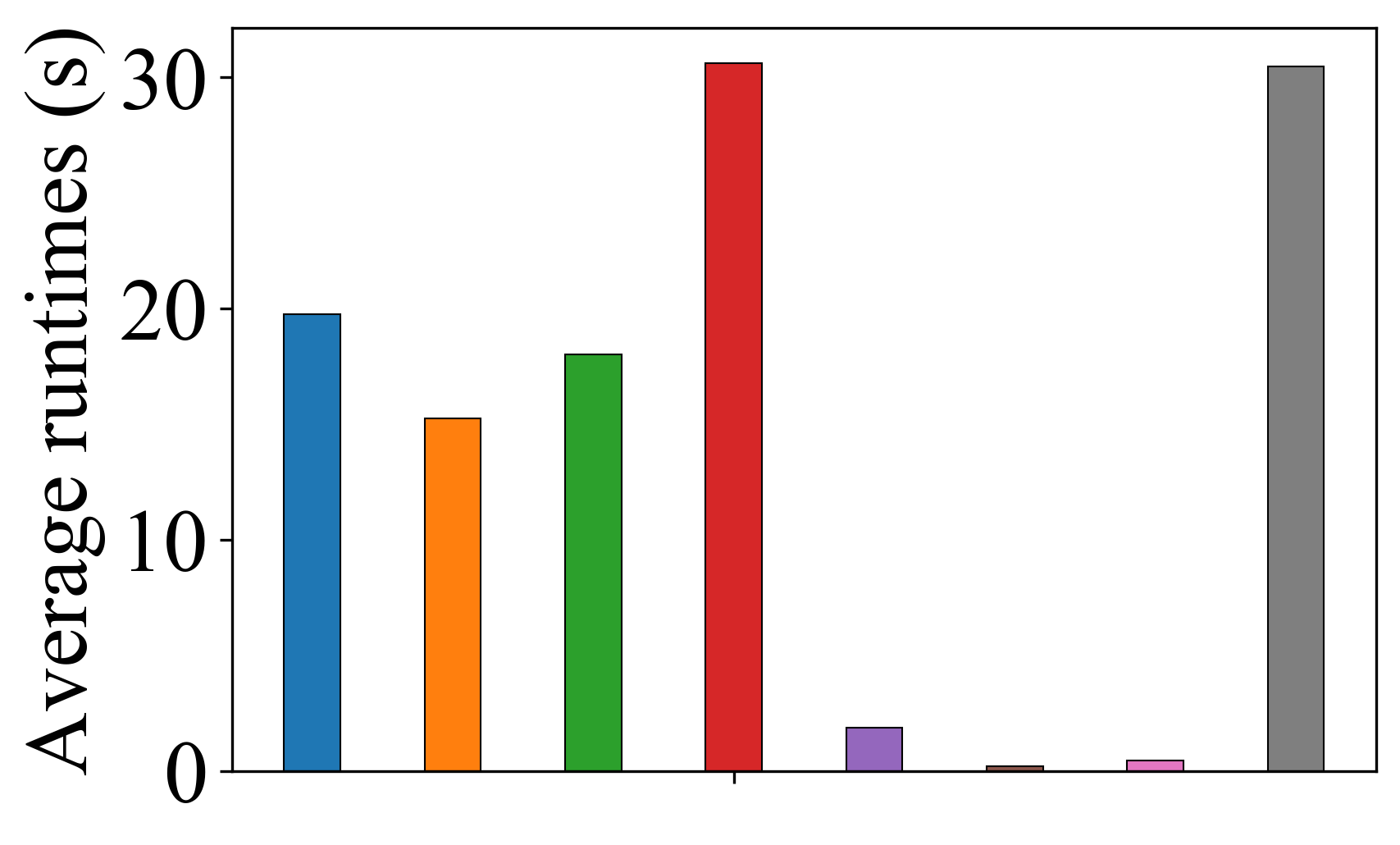

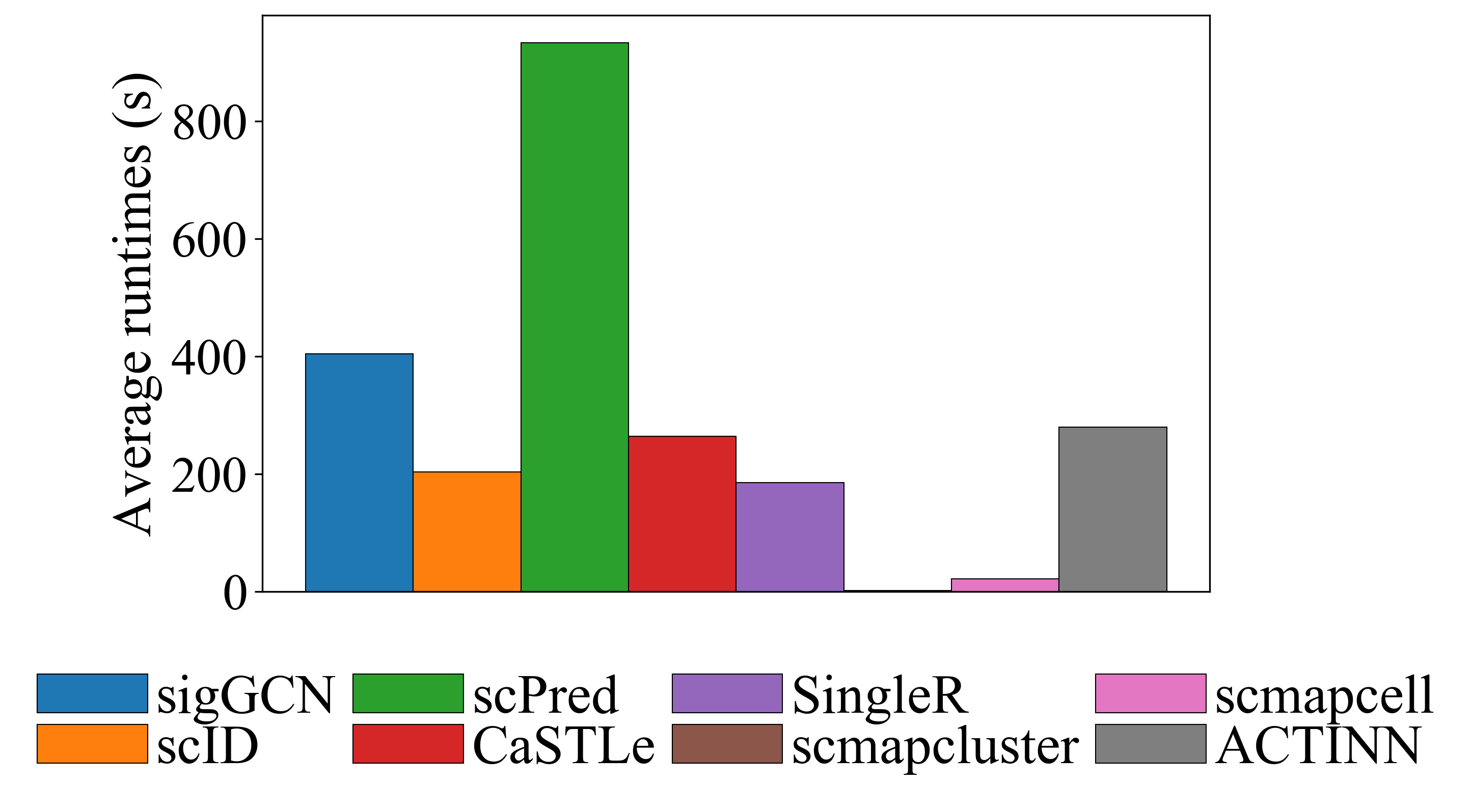

Figure S15: Average runtimes of the methods using (a) Zheng68K (b) Baron Human (c) Muraro (d) Segerstolpe (e) Baron Mouse (f) Xin dataset.
